# TNF-mediated neuroinflammation is linked to neuronal necroptosis in Alzheimer’s disease hippocampus

**DOI:** 10.1101/2021.07.09.451781

**Authors:** Anusha Jayaraman, Thein Than Htike, Rachel James, Carmen Picon, Richard Reynolds

## Abstract

The pathogenetic mechanisms underlying neuronal death and dysfunction in Alzheimer’s disease (AD) remain unclear. However, chronic neuroinflammation has been implicated in stimulating or exacerbating neuronal damage. The tumor necrosis factor (TNF) superfamily of cytokines are involved in many systemic chronic inflammatory and degenerative conditions and are amongst the key mediators of neuroinflammation. TNF binds to the TNFR1 and TNFR2 receptors to activate diverse cellular responses that can be either neuroprotective or neurodegenerative. In particular, TNF can induce programmed necrosis or necroptosis in an inflammatory environment. Although activation of necroptosis has recently been demonstrated in the AD brain, its significance in AD neuron loss and the role of TNF signaling is unclear. We demonstrate an increase in expression of multiple proteins in the TNF/TNF receptor-1-mediated necroptosis pathway in the AD post-mortem brain, as indicated by the phosphorylation of RIPK3 and MLKL, predominantly observed in the CA1 pyramidal neurons. The density of phosphoRIPK3+ and phosphoMLKL+ neurons correlated inversely with total neuron density and showed significant sexual dimorphism within the AD cohort. In addition, apoptotic signaling was not significantly activated in the AD brain compared to the control brain. Exposure of human iPSC-derived glutamatergic neurons to TNF increased necroptotic cell death when apoptosis was inhibited, which was significantly reversed by small molecule inhibitors of RIPK1, RIPK3, and MLKL. In the post-mortem AD brain and in human iPSC neurons to TNF, we show evidence of altered expression of proteins of the ESCRT III complex, which has been recently suggested as an antagonist of necroptosis and a possible mechanism by which cells can survive after necroptosis has been triggered. Taken together, our results suggest that neuronal loss in AD is due to TNF-mediated necroptosis rather than apoptosis, which is amenable to therapeutic intervention at several points in the signaling pathway.

## Introduction

In recent years, accumulating evidence suggests that the longstanding amyloid cascade hypothesis cannot sufficiently explain many aspects of Alzheimer’s disease (AD) pathogenesis, bringing the exploration of other possible underlying mechanisms to the forefront [30]. Neuroinflammation is suggested to play a major role in ongoing neurodegeneration in AD, as demonstrated by increased inflammatory markers in patients with AD, the discovery of AD risk genes associated with innate immunity [21], and the prominent presence of dysregulated immune pathways in the genomic and transcriptomics analyses of AD tissues [64]. One of the key mediators of inflammation in many systemic chronic inflammatory and degenerative conditions is the soluble form of tumor necrosis factor (TNF), acting primarily through its binding to TNF receptor 1 (TNFR1) [12], and there is increasing evidence that TNF signaling is also implicated in multiple CNS degenerative conditions [63]. TNF binding to the death domain containing TNFR1, and subsequent formation of complex 1, can lead to either induction of NFkB signaling or cytotoxicity via apoptosis or necroptosis following formation of complex 2. Apoptosis is triggered when the receptor interacting kinase 1 (RIPK1) is deubiquitinated by CYLD protein and then cleaved by activated caspase 8 [36, 38]. However, under chronic inflammatory conditions characterized by down-regulated caspase 8 activation, RIPK1 interacts with RIPK3 to induce autophosphorylation, leading to the recruitment and phosphorylation of pseudokinase mixed lineage kinase domain-like (MLKL), which together form the necrosome. pMLKL oligomers then translocate to the plasma membrane and disrupt it, leading to necroptotic cell death [53, 57].

Apoptotic neurons are rarely detected in the AD brain [25], whereas necroptosis has been recently implicated in several neurodegenerative and neuroinflammatory disorders, including multiple sclerosis [39, 42], AD [5, 28] and amyotrophic lateral sclerosis [44], but the triggering and signaling mechanisms underlying necroptotic cell death in these conditions are still unclear. There are a number of known triggers of necroptotic signaling in non-neuronal cells, of which TNF is the most studied [68]. Although TNF is suggested to be a key mediator of chronic inflammation and cytotoxicity in many neurodegenerative and neuroinflammatory disorders [63], few comprehensive human tissue studies have been undertaken to investigate the signaling pathways concerned and the cellular specificity. Levels of TNF in the cerebrospinal fluid (CSF) have been shown to be increased in AD at an early stage [23, 54, 55] and TNF and TNFR1 protein levels have been reported to be increased in early-stage AD post-mortem brains [67]. Treatment of primary microglial cultures with Aβ has been shown to result in high levels of TNF release from these cells [8]. Chronic neuron-specific expression of TNF in 3xTg-AD mice has been shown to result in inflammation-driven neuronal death in this AD mouse model [24]. In addition, a single nucleotide polymorphism in the TNF gene, G308A, has been reported to have a significant association with susceptibility to AD in certain populations, while being protective in others [59]. Our previous studies on neurodegeneration in the MS brain indicate that, in this prototypical neuroinflammatory condition, TNF/TNFR1 interaction and downstream activation of the RIPK1/RIPK3/MLKL kinase cascade is the most likely cause of neuronal necroptosis [42]. Here we show the activation of the TNFR1-mediated necroptosis pathway in hippocampal CA1-2 neurons and a concomitant downregulation of apoptotic signaling in a cohort of post-mortem AD brains. In addition, we report a dysregulation in the ESCRTIII pathway, which has been suggested to be involved in the rescue of cells from necroptosis in non-CNS systems [16]. Treatment of human iPSC-derived glutamatergic neurons with TNF led to necroptotic cell death when apoptosis was inhibited. Small molecule inhibitors of RIPK1, RIPK3 and MLKL protected these neurons against TNF-mediated cell death, suggesting new potential therapeutic avenues against neurodegeneration in the AD brain.

## Materials and Methods

### Tissue samples

The AD post-mortem tissues for this study were obtained from the Multiple Sclerosis and Parkinson’s Tissue Bank at Imperial College London and the South West Dementia Brain Bank, University of Bristol. Snap frozen tissue blocks of anterior hippocampus and entorhinal cortex were obtained from 30 AD cases (12 males, 18 females; median age of death = 85.5 years, range = 57–99 years; Braak stages III–VI) and 11 control cases (6 males, 5 females; median age of death = 83 years, range 63–95 years). Formalin fixed paraffin embedded (FFPE) sections were obtained for a subset of cases and controls (n=5 per group) from the Multiple Sclerosis and Parkinson’s Tissue Bank at Imperial College London. Fully informed consent was obtained for the post-mortem donation under ethical approval by the National Research Ethics Committee (08/MRE09/31 and NHS REC No 18/SW/0029). The project was approved by the Nanyang Technological University Institutional Review Board (IRB-2018-09-052). The demographic data and neuropathological features of the AD cases provided by both brain banks were determined in accordance with the standardised criteria of the UK MRC Brain Bank Network and are shown in Supplemental Table 1.

### Immunohistochemistry and immunofluorescence

For immunofluorescence analysis, 10 μm cryosections containing the CA1 region were cut from the snap frozen hippocampal blocks. For immunohistochemistry, snap frozen tissue sections were fixed with formalin for 30 min followed by ice cold methanol for 10 min. Slides were blocked with 10% normal goat or horse serum followed by overnight incubation with primary antibody at 4°C and 30 min incubation with ImmPRESS HRP-conjugated secondary antibodies (Vector Laboratories) at room temperature. Slides were visualized with ImmPACT-DAB (Vector Laboratories) as the chromogen. Sections were counterstained with hematoxylin and DPX (Sigma-Aldrich) mounted. Dual color IHC was performed sequentially. Detection of the primary antibody with ImmPACT-DAB was followed by incubation with the second primary antibody, which was detected using the ABC-alkaline phosphatase detection system (Vector Laboratories), using Vector blue as the substrate. When using human paraffin embedded sections for IHC, the sections were deparaffinized and subjected to heat-induced epitope retrieval using citrate buffer before following the same steps as for snap-frozen tissue. For immunofluorescence, 10 μm human tissue cryosections were fixed with ice cold methanol, blocked with 10% normal goat or horse serum, and incubated overnight with primary antibodies. Sections were then incubated with the appropriate secondary antibody conjugated to a fluorochrome and mounted with Vectashield^®^ Vibrance^™^ Antifade Mounting medium with DAPI (Vector Laboratories) for nuclei counterstaining. Individual antibody details are listed in Suppl. Table 2.

### Protein extraction

Hippocampal grey matter from snap-frozen human AD and control samples was dissected by carefully scoring the tissue block with a fine scalpel and subsequent isolation of the tissue by cryosectioning in a Leica cryostat. The resultant tissue samples were homogenized in RIPA buffer (ab156034, Abcam) containing protease and phosphatase inhibitors (ab201119, Abcam) and incubated on ice for 20 min at 4°C. The protein extract was centrifuged at 14800 g for 15 min at 4°C. The resulting supernatant was taken as the RIPA soluble fraction. Pellets were washed in TBS and homogenized in 6 M urea/5% SDS for 30 min at room temperature (RT) and centrifuged at 14800 g for 10 min at 4°C. The resultant supernatant was the urea fraction. The samples were stored at −80 °C until further use. To extract proteins in native condition, tissue was homogenized in TBS containing 0.1% of Triton X-100, incubated for 10 min at RT, followed by centrifugation at 16000 g for 10 min.

### Western blotting

Briefly, protein concentration was quantified using a Sigma-Aldrich BCA protein assay kit (Merck). Subsequently, 10-50 μg of protein was loaded onto 4-15% Mini-PROTEAN^®^ TGX Stain-Free^™^ Protein Gels (Bio-Rad) and transferred to polyvinylidene difluoride (PVDF) membranes for 40-60 min. The membranes were incubated for 1 h in 5% BSA at room temperature and incubated overnight at 4°C with the appropriate primary antibodies. The blots were then washed 3x with TBS-T for 10 min each and incubated with the horseradish peroxidase (HRP) conjugated specific secondary antibodies (1:20,000, Bio-Rad) for 1 h at RT. The blots were then washed with TBS-T, and imaged/quantified using a Bio-Rad ChemiDoc^™^ MP Imaging System and normalized with GAPDH expression. For the detection of MLKL oligomers, proteins were run on 7.5% Mini-PROTEAN^®^ TGX Stain-Free^™^ Protein Gel (Bio-Rad) in non-reducing conditions. Individual antibodies are listed in Suppl. Table 2.

### RNA extraction and RT-PCR

For RNA extraction, grey matter regions in each brain tissue block were demarcated with a scalpel and then cryosectioned as above. The grey matter tissue sections were then homogenized and processed for total RNA extraction using PureLink^™^ RNA Mini kit (Life Technologies Corporation) as per the manufacturer’s protocol. Purified total RNA (100 ng) was used from each sample for One-step reverse transcriptase quantitative polymerase chain reaction (RT-qPCR) using the iTaq^™^ Universal SYBR^®^ Green One-Step kit (Bio-Rad) in the StepOnePlus^™^ Real Time PCR system (Applied Biosystems). All primers in the study were commercially purchased PrimePCR^™^ SYBR^®^ Green assay primers (Bio-Rad). For each sample, reactions were set up in triplicate with the following cycling protocol: 50°C for 10 min, 95°C for 1 min, 40 cycles with a 3-step protocol (95°C for 15 sec, 60°C for 1 min), and a final melting curve analysis with a ramp from 65-95°C. Relative quantification of mRNA levels from various treated samples was determined by the comparative Ct method [48] after normalizing with the corresponding *xpnpep1* levels [9] from the samples.

### Immunoprecipitation

Samples from AD and control brain grey matter were homogenized in RIPA buffer containing protease and phosphatase inhibitors. Immunoprecipitation was carried out using the Dynabeads^™^ Protein G Immunoprecipitation Kit (ThermoFisher Scientific) according to the manufacturer’s protocol. Briefly, 10 μg of anti-MLKL antibody was incubated for 10 min with 50 μL of Dynabeads Protein G while rotating. The beads were then washed and incubated with control and AD lysate samples overnight at 4°C. The beads were then washed three times with washing buffer and resuspended in the elution buffer mixed with 2X sample buffer (Bio-Rad). The beads were boiled at 95°C for 5 min and placed on ice. Bead-free supernatant samples were loaded onto a 4--15% Mini-PROTEAN^®^ TGX Stain-Free^™^ Protein Gels. Gels were transferred to a PVDF membrane and incubated overnight at 4°C with anti-pRIPK3 followed by 1 h incubation with anti-rabbit IgG-HRP secondary antibody at RT. Blots were then washed and developed with Amersham^™^ ECL^™^ Select WB Detection Reagent. Individual antibodies are listed in Suppl. Table 2.

### IPSC glutamatergic neuron culture

Human ioNEURONS/glut glutamatergic neurons (ab259259, Abcam) were seeded at 2×10^5^cell/well on 96 well plates (Biolab) coated with Geltrex in Neurobasal media supplemented with B27 Plus, GlutaMax^™^, penicillin/streptomycin, NT3 and BDNF (Life Technologies) as per the manufacturer’s protocol. The cultures were maintained in a humidified atmosphere of 5% CO_2_ in air at 37°C. All experiments were performed at 9-11 days post-plating. Treatment with TNF (ThermoFisher), SMAC mimetics (Tocris) and caspase inhibitor Z-VAD-FMK (Abcam) (TSZ treatment) was performed at the concentrations indicated in the text. Necroptosis inhibitors GSK-547 (SelleckChem), GSK-872 (Abcam) and necrosulfonamide (Abcam) were added to the cultures 1 h before the treatment with TSZ at concentrations indicated in the text.

### LDH assay

Cell cytotoxicity was determined by measuring the lactate dehydrogenase (LDH) release (ab65393, LDH-Cytotoxicity Assay kit, Abcam) from iPSC ioNEURONS/glut under different treatment conditions. Ten μl supernatant samples from the neuronal cultures under different conditions were collected and processed following the manufacturer’s protocol. The LDH released was determined by measuring the absorbance of the samples with a Synergy H1 microplate reader (BioTek) at 450 nm.

### Reactive oxygen species assay

A reactive oxygen species (ROS) assay was carried out using the Cellular ROS assay kit (ab133854, Abcam) according to the manufacturer’s protocol for adherent cells. Cells under different treatment conditions were incubated with the diluted 2’,7’-dichlorodihydrofluorescein diacetate (DCFDA) solution for 45 min at 37°C in the dark. The solution was removed, and the cells were washed once with PBS. The culture plate was immediately measured on a fluorescence plate reader (Synergy H1, BioTek) at Ex/Em = 485/535 nm in endpoint mode. DCFDA is taken up by the cells, where in the presence of ROS, it is oxidized and converted to 2’,7’-dichlorofluorescein (DCF), which is highly fluorescent. ROS was quantified by measuring the fluorescence of DCF.

### Image acquisition and analysis

Immunohistochemistry slides were digitized by whole slide scanning using a Zeiss Axio Scan Z1 scanner. Image files were handled using QuPath v0.2.0. open-source software for analyses. HuC/D, MHC-II, pRIPK3 and pMLKL cell densities were measured with the “Positive Cell Detection of a Region of Interest” tool in QuPath. The total number of cells for all the experiments is given as total cells/mm^2^. Immunofluorescence images were captured using an Axio Observer 7 Inverted fluorescence microscope (Zeiss) and processed with Zen 2 (Blue edition) software for analyses.

### Statistical analysis

All human post-mortem data was assumed to be sampled from a non-Gaussian distribution and non-parametric analysis methods applied. The difference between two groups was compared using the unpaired Mann-Whitney test. Spearman correlation was used to test for associations between groups and the Spearman r and p values reported in each instance. One-way ANOVA with Bonferroni’s post-hoc correction was performed for multiple comparisons. Paired t-tests were used for comparing two groups in culture experiments. A two-sided p-value <0.05 was considered significant.

## Results

### Necroptotic signaling is highly activated in AD hippocampus

In order to demonstrate the activation of necroptotic signaling in hippocampal neurons in the AD brain, we studied the detailed changes in this pathway at both mRNA and protein levels in brain tissues from 20-30 AD patients and 10 non-neurological controls. There was a 2-3-fold increase in the mRNA expression levels for *RIPK1* (mean±SEM AD: 0.8±0.18, Ctrl: 0.2±0.06; p = 0.0152), *RIPK3* (AD: 1.3±0.14, Ctrl: 0.5±0.15; p = 0.0007) and *MLKL* (AD: 1.33±0.15, Ctrl: 0.68±0.12; p = 0.0057) in AD cases (Fig. 1a). Although there were some differences in the levels of RIPK1, RIPK3, and MLKL protein in the UREA soluble fraction, determined by western blotting, these were not significantly different between controls and AD, mainly due to the high degree of heterogeneity in the AD cases (Fig. 1b, c). In contrast, the activated pRIPK3 protein (AD: 4.04±0.54; Ctrl: 1.93±0.36; p = 0.0101) and pMLKL protein (AD: 1.96±0.21; Ctrl: 0.94±0.1; p = 0.0024) levels were significantly upregulated in the hippocampal grey matter of AD cases compared to controls (Fig. 1b, c). Using non-reducing conditions to probe the oligomeric state of MLKL, we found the almost exclusive presence of MLKL oligomers of approximately 250 kDa in the AD hippocampus, most likely representing tetramers (Fig. 1d). Immunohistochemical (IHC) localization of the expression of pRIPK3 and pMLKL, demonstrated pMLKL+ (Fig 2a) and pRIPK3+ (Fig 2b) staining in the AD hippocampus, but not in the controls. The staining was characteristic of the granulovacuolar degeneration (GVD) bodies in pyramidal neurons and 9.4%±1.3% and 14.45%±2.55% of total neurons in the CA1 and CA2 regions of AD cases expressed pMLKL and pRIPK3 respectively, compared to 0.04±0.03% and 0.003%±0.002%, in controls (Fig. 2a, b; Suppl. Fig 1a). There was no colocalization of pMLKL immunostaining with astrocytes or microglia (Suppl Fig 1b). In addition, we also observed significantly higher numbers of pMLKL (2-fold) and pRIPK3 (2.5-fold) positive neurons in female AD brains when compared to male AD brains (Suppl. Fig 1c). Therefore, our data suggests activation of the necroptosis pathway in CA1-2 hippocampal pyramidal neurons in AD brain with some evidence of sexual dimorphism.

**Fig. 1.**
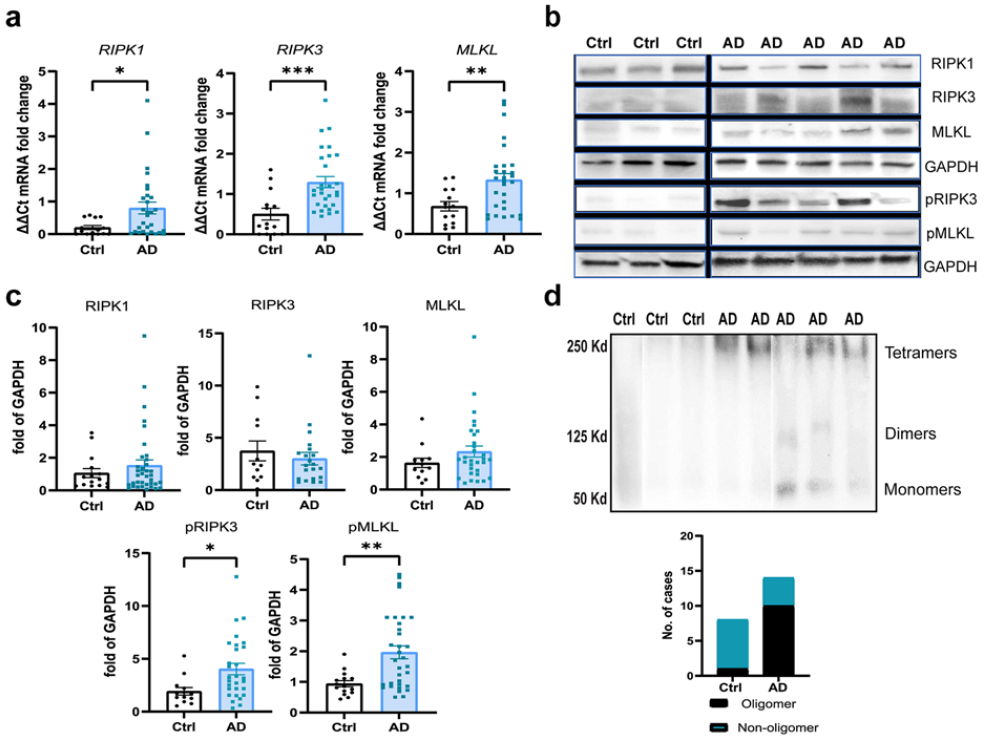
Activation of necroptosis signaling in AD hippocampus. **a)** Analysis of mRNA levels of the RIPK1, RIPK3, and MLKL genes in the hippocampus grey matter in AD cases (n=30) and controls (n=14). **b)** Representative western blots of RIPK1, RIPK3/pRIPK3, and MLKL/pMLKL. Full blots are shown in suppl Fig. 6. **c)** Quantification of RIPK1, RIPK3, MLKL, pRIPK3, and pMLKL protein levels in AD (n=20) and control cases (n=10), normalized to GAPDH. **d)** Western blot analysis of MLKL from hippocampal grey matter tissue lysate extracted in native conditions, showing the presence of oligomers. Quantification of the MLKL oligomers (~250 kDa) in the AD (n=14) and control (n=8) cases. For two group comparisons, the Mann-Whitney test was used. Data are represented as mean ± SEM, *p<0.05, **p<0.01, ***p<0.001, ****p<0.0001.

**Fig. 2.**
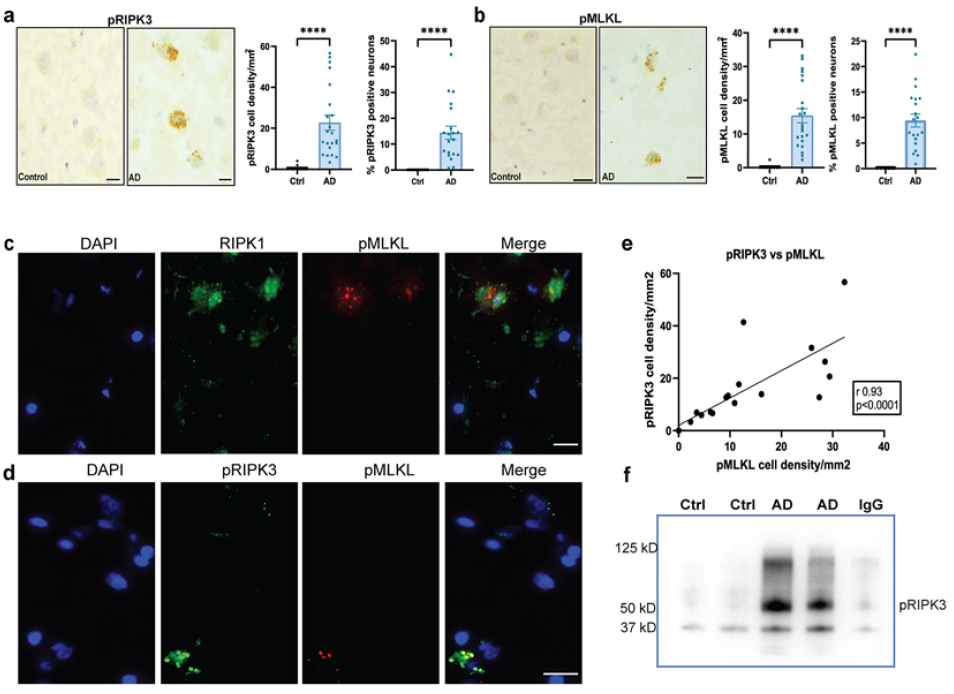
Necrosome formation in the AD hippocampus. **a)** Representative images of pRIPK3 IHC in the hippocampal CA1 region from a control and AD case (scale bar = 20μm) and the quantification of pRIPK3+ cell density and percentage of pRIPK3+ neurons in AD (n=22) and control (n=10) cases. **b)** Representative images of pMLKL IHC in the hippocampal CA1 region from a control and AD case (scale bar = 20μm) and the quantification of pMLKL+ cell density and percentage of pMLKL+ neurons in AD (n=22) and control (n=10) cases. Data are represented as mean ± SEM, ****p<0.0001. **c)** Representative immunofluorescence image of RIPK1 (green) and pMLKL (red) with DAPI+ cell nuclei (blue) in CA1 pyramidal neurons of an AD case (scale bar = 20μm). **d)** Representative immunofluorescence image of pRIPK3 (green) and pMLKL (red) with DAPI+ cell nuclei (blue) in CA1 pyramidal neurons of an AD case (scale bar = 20μm). **e)** Correlation analysis between pRIPK3 and pMLKL cell densities in AD cases (r = 0.93, p<0.0001). **f)** Co-immunoprecipitation analysis of pRIPK3 with MLKL in the hippocampal grey matter lysates from 2 AD and 2 control cases. Correlation analysis by Spearman comparison.

### Necrosome formation in the AD hippocampus

Previous studies have shown that the necrosome complex is formed when RIPK1 binds to and phosphorylates RIPK3, which further binds to and phosphorylates MLKL, leading to the oligomerization and migration of pMLKL to the plasma membrane [57]. In this regard, the presence of pMLKL is an indication of activation of the necroptosis pathway. Therefore, we have considered this a marker for necroptosis throughout the study. We observed co-expression of pMLKL along with RIPK1 (Fig 2c) and pRIPK3 (Fig 2d) in the same neurons in the AD hippocampus. Interestingly, all pMLKL+ neurons were also pRIPK3+, showing colocalization at the subcellular level (Fig 2d), but some pRIPK3+ pMLKL-neurons (arrowheads) were also present. This suggests that these neurons were at different stages of the necroptosis pathway. We also found a strong correlation between pRIPK3+ and pMLKL+ cell densities in the AD cases (Fig 2e). Co-immunoprecipitation of pRIPK3 with MLKL could be demonstrated in the AD hippocampus but not in the controls (Fig 2f), which further provides evidence for the activation of the necroptosis pathway and formation of the necrosome complex in the AD hippocampus.

### Association between pMLKL+ neurons and AD pathology

To investigate if there was any difference in neuron numbers between AD cases and control cases, we stained the hippocampal sections for the neuronal marker, HuC/HuD and calculated the neuron density per mm^2^ in the CA1-2 area. We observed that there was a significant 45% neuron loss in the AD hippocampus (99.95±10.78 neurons/mm^2^) when compared with the control samples (182.3±17.6 neurons/mm^2^; p = 0.0006) (Fig 3a). In addition, we observed a strong negative correlation between total neuron density and pMLKL+ cell density (r −0.73, p < 0.0001) and between total neuron density and pRIPK3+ cell density (r −0.63, p<0.01)(Suppl Fig 1d). A previous study on necroptosis in AD had suggested an association between necroptosis and Tau pathology, but not Aβ pathology [28]. In our AD cohort, we found that there was a positive correlation between pMLKL cell density and pTau levels, but not with Aβ levels (Suppl Fig 2a, b). Similarly, we found a significant positive correlation between pRIPK3 and pTau levels, but not between pRIPK3 and Aβ levels (Suppl. Fig 2e, f). However, pTau and Aβ Braak stages showed significant correlations with pMLKL+ cell density (Suppl Fig 2c, d), but not pRIPK3 cell density (Suppl Fig 2g, h). Immunostaining showed co-expression of pTau+ neurofibrillary tangles and pMLKL in some neurons (Fig 3b). In contrast, there was no overlap in the expression of Aβ and pMLKL within the same neurons (Fig 3c). However, pMLKL+ neurons were often found in close proximity to Aβ plaques (Fig 3c).

**Fig. 3.**
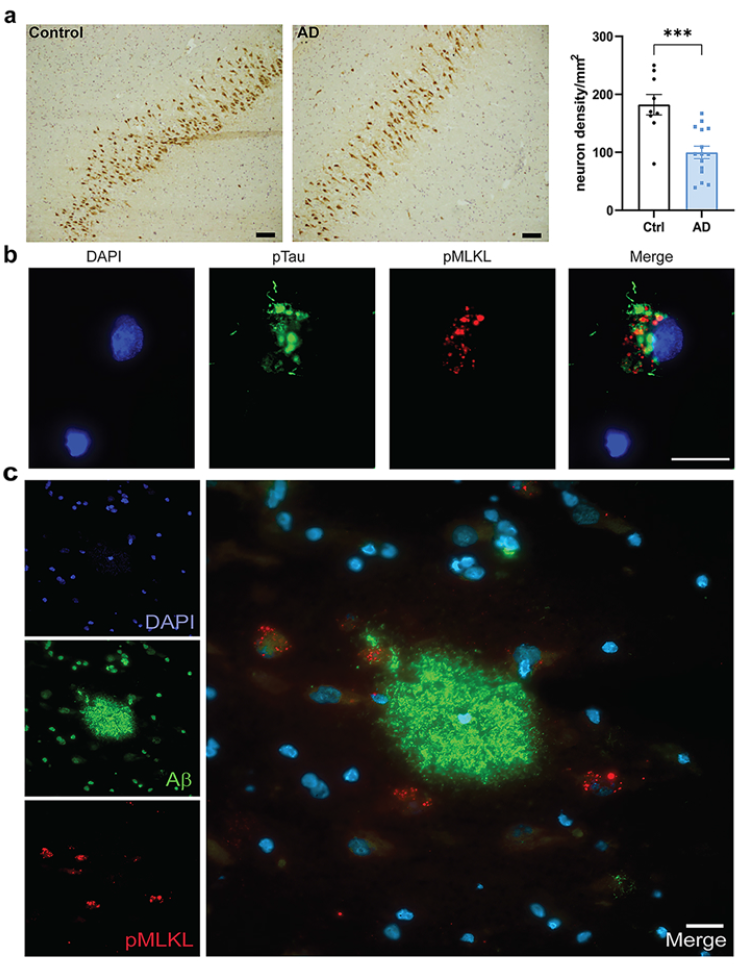
Association of pMLKL with AD pathology. **a)** Representative images of CA1 pyramidal neurons stained with HuC/HuD antibody in the control and AD brain sections (scale bar = 100μm); Quantification of the neuron density per mm^2^ in the control and AD hippocampal CA1 region. Data are represented as mean ± SEM, ***p<0.001. **b)** Representative immunofluorescence image of pTau (green) and pMLKL (red) with DAPI+ cell nuclei in CA1 region of AD hippocampus. **c)** Representative immunofluorescence image of Aβ (green) and pMLKL (red) with DAPI+ cell nuclei in CA1 region of AD hippocampus. Scale bar = 20μm.

The AD brain sections showed a significantly increased presence of HLA-DR+ microglia compared to the control sections, as measured by the mean HLA-DR intensity (Ctrl: 2.08±0.1, AD:3.55±0.32; p = 0.0035), suggesting significant microglial activation in the AD tissue (Fig 4a). The activated HLA-DR+ microglia in the AD brains were present both in the white matter and grey matter, with morphologies ranging from moderately hypertrophic cell bodies to highly activated enlarged cell bodies. We also observed increased numbers of CD8+ T cells in the AD cases compared to the control cases, which were predominantly present in the perivascular spaces and meninges but were also observed to a lesser extent within the parenchyma, especially in the AD hippocampus (Fig 4b). In the AD cases, the activated HLA-DR+ microglia and the CD8+ T cells were observed in the proximity of pMLKL+ neurons (Fig 4c, d). Moreover, the mean intensity of HLA-DR staining and the overall CD8+ T cell density correlated significantly with the pMLKL+ neuron density in the AD hippocampus (Fig 4c, d). Thus, our data suggests several factors that could contribute to a chronic inflammatory microenvironment around specific CA1 neurons, increasing their susceptibility to necroptosis.

**Fig. 4.**
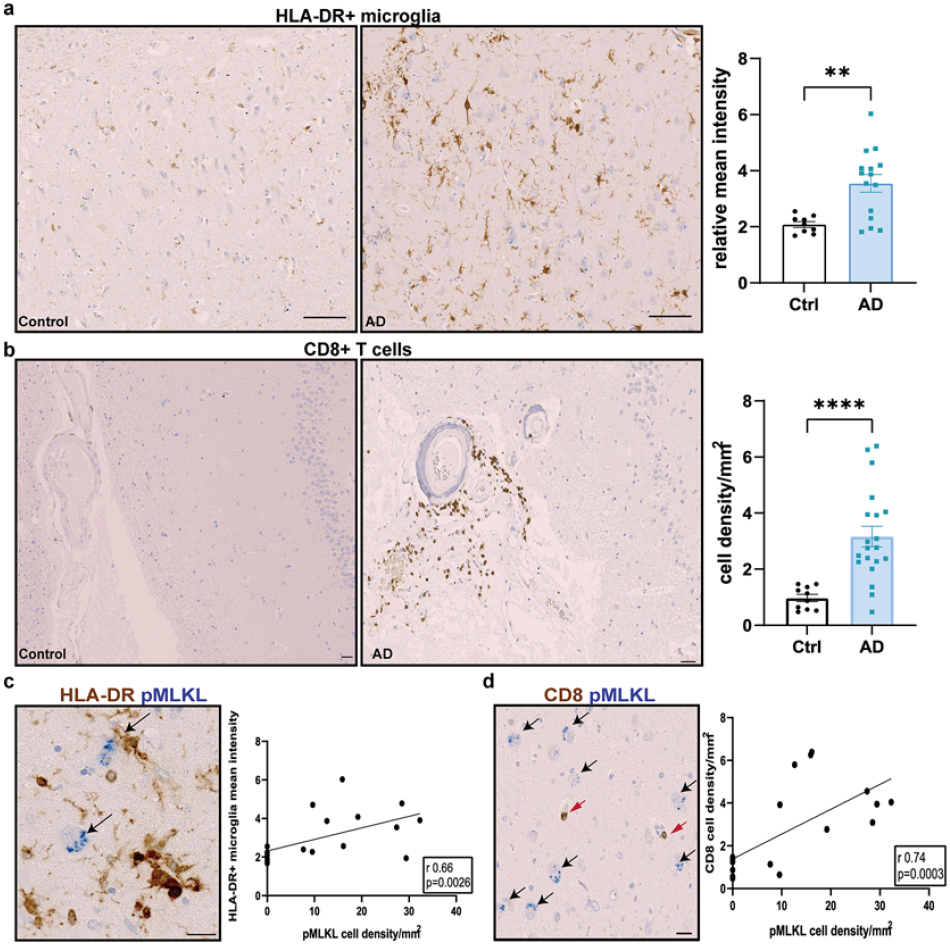
Representative IHC and quantification of HLA-DR positive microglia in control and AD brain sections (scale bar = 100μm). Data are represented as mean ± SEM, **p<0.01. **b)** Representative IHC showing CD8+ T cells (brown) in the control (left) versus AD (right) tissues, along with the presence of CD8+ T cells in close proximity to pMLKL+ neurons in the CA1 region of the AD hippocampus (scale bar = 100μm). **c)** Representative image showing pMLKL+ neurons (black arrows) in close proximity to HLA-DR+ microglia (scale bar = 20μm), along with correlation analysis between mean HLA-DR intensity and pMLKL+ neuron density (r = 0.66, p = 0.0026). **d)** Representative image showing pMLKL+ neurons (black arrows) in close proximity to CD8+ T cells (red arrows) (scale bar = 20μm), along with correlation analysis between CD8+ T cell density and pMLKL+ neuron density (r = 0.74, p = 0.0003). Correlation analysis by Spearman comparison.

### TNFR1/RIPK1 signaling is upregulated in neurons in the AD hippocampus

The initiating factors underlying necroptotic cell death in neurodegenerative diseases are still unclear. In non-neuronal model systems, TNF is the best-known trigger of necroptosis [68]. We have previously demonstrated the presence of TNF-mediated necroptosis signaling in MS [42]. Therefore, we investigated the link between TNF signaling and necroptosis in AD brain by determining the expression levels of TNF receptors 1 and 2, and a key initial regulator Fas associated death receptor (FADD). A significant upregulation of *TNFR1* (Ctrl: 1.74±0.25, AD:3.75±0.38; p = 0.0002) and *TNFR2* (Ctrl:0.17±0.04, AD: 0.51±0.05; p <0.0001) gene expression, but no change in *FADD* gene expression, was observed in the AD brain compared to controls (Fig 5a). Western blot analysis showed a significant increase in both soluble TNF (17 kD; Ctrl: 0.7±0.23, AD:2.42±0.52; p = 0.0247), the membrane-bound form of TNF (35 kD; Ctrl: 1.33±0.31, AD: 3.46±0.65; p = 0.0421) and FADD (Ctrl: 0.67±0.13, AD: 1.73±0.35, p = 0.0473) levels. There was a clear trend towards an increase in TNFR1 protein levels (Ctrl: 1.08±0.31, AD: 2.18±0.31; p = 0.063), which did not reach significance due to substantial heterogeneity between cases, but no difference in TNFR2 levels (Fig 5b, c). As reported previously (Yuan et al., 2019), we observed a low level of TNFR1 expression in all CA1 neurons in both control and AD cases. However, triple immunofluorescence revealed increased TNFR1 expression specifically in neurons in the AD hippocampus that were pRIPK3+ or pMLKL+ (Fig 5d, e). Together with the demonstration of upregulation and localisation of the downstream components of the necroptosis pathway (Figs 1–2), these results strongly suggest a TNFR1-mediated activation of necroptosis in these neurons in the AD hippocampus.

**Fig. 5.**
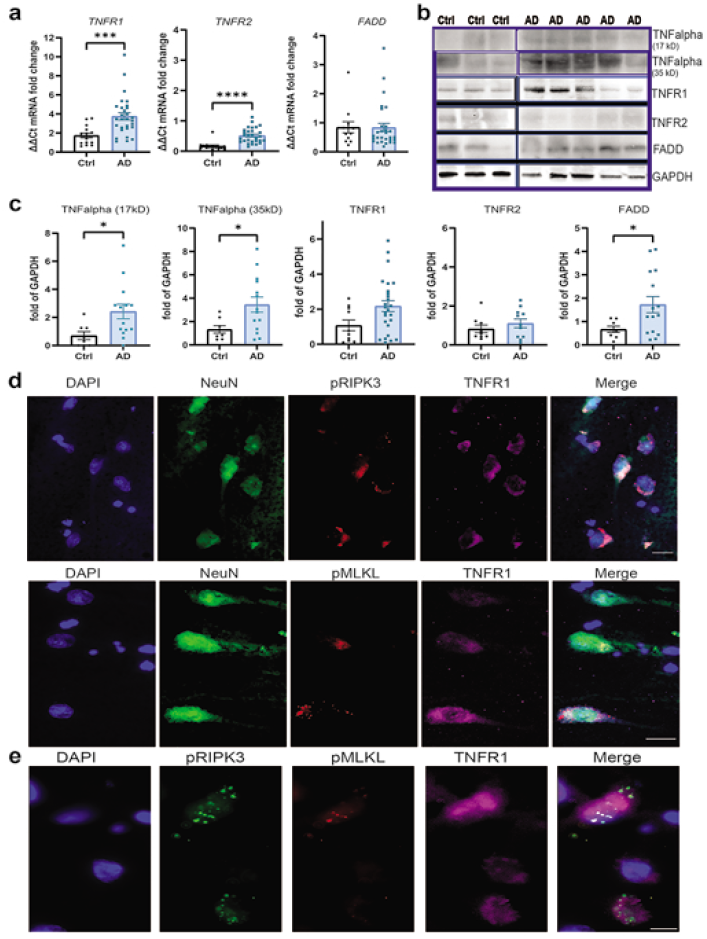
TNFR1-mediated necroptosis signaling is activated in the CA1 neurons in AD hippocampus. **a)** Analysis of mRNA levels of the TNFR1, TNFR2, and FADD genes in the hippocampus grey matter in AD cases (n=30) and controls (n=14). **b,c)** Representative western blots of the soluble and membrane bound forms of TNF and TNFR1, TNFR2, and FADD (full blots are shown in suppl Fig. 6) and their quantification in AD (n≥10) and control cases (n≥5), normalized to GAPDH. Data are represented as mean ± SEM, *p<0.05, ***p<0.001. **d)** Representative immunofluorescence image of pRIPK3 (red, top panel) and pMLKL (red, bottom panel) and TNFR1 (pink) in NeuN+ (green) CA1 neurons with DAPI+ cell nuclei (blue) in an AD case (scale bar = 20μm). **e)** Representative immunofluorescence image of pRIPK3 (green), pMLKL (red) and TNFR1 (pink) with DAPI+ cell nuclei (blue) in an AD case (scale bar = 20μm).

### Apoptosis signaling is not activated in the AD hippocampus

To determine the extent of apoptosis signaling in the AD hippocampus, we next studied the activation of caspase dependent apoptosis. RT-PCR analysis showed a significant decrease in *CASP3, cIAP1* and *cFLAR* expression levels in the AD brain (Fig 6a). Caspase-dependent apoptosis is initiated by proteolytic cleavage of caspase-8 (55 kD) into the p18 and p43 subunits [22]. The protein levels of the cleaved active p18 subunit were not significantly different from that in control samples (Fig. 6b). However, the downstream cleaved caspase-3 levels were significantly lower in the AD brain when compared to control samples (Fig 6b). These results, along with lower gene expression levels of *cFLAR* and *cIAP1*, indicate that caspase-dependent apoptosis is not significantly activated in neurons in AD, which suggests that extrinsic activation of the apoptotic pathway is not the main mechanism driving cell death in neurons in AD.

**Fig. 6.**
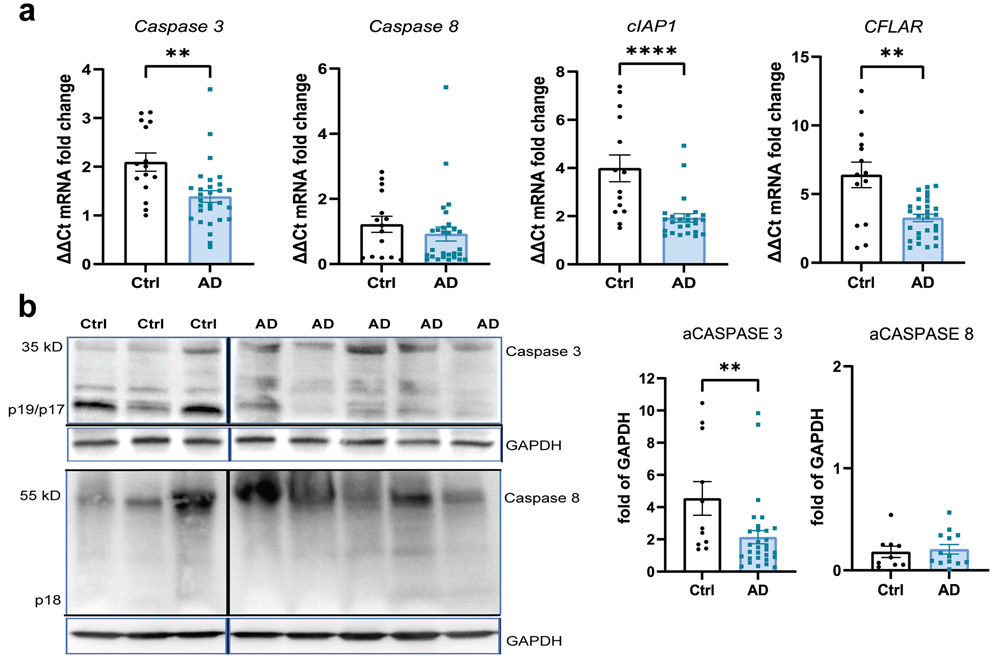
Apoptosis signaling is downregulated in AD hippocampus. **a)** Analysis of mRNA levels of the Caspase 3, Caspase 8, cIAP1, and CFLAR genes in the hippocampus grey matter in AD cases (n=30) and controls (n=14). **b)** Representative western blots of full-length and cleaved caspase 3 and caspase 8. Full blots are shown in suppl Fig. 6. Quantification of cleaved caspase 3 and caspase 8 protein levels in AD (n≥10) and control cases (n≥6), normalized to GAPDH. Data are represented as mean ± SEM, **p<0.01, ***p<0.001, ****p<0.0001.

### ESCRT III pathway proteins are altered in AD hippocampus

The endosomal sorting complexes required for transport group III (ESCRT III) proteins, have been recently implicated as a counterbalance for necroptosis and a possible inherent mechanism to prevent or delay cell death after necroptosis has been triggered. Conversely, silencing of ESCRT-III can induce necroptosis in cells [16]. CHMP2B, a component of the ESCRT III complex, is also a marker for GVD bodies [61]. Therefore, we investigated if key components of the ESCRT-III pathway showed altered expression in the AD hippocampus. Gene expression analysis of the essential components of the ESCRT III complex (*CHMP2a, CHMP2b, VPS24/CHMP3, CHMP4b and CHMP6*) and *VPS4b*, which is important for recycling the ESCRT-III components, showed that *VPS4b* and *CHMP3* had significantly higher expression in the AD brains compared to the controls (Fig 7a, Suppl. Fig 3a). At the protein level, however, only Chmp3 showed a significant increase in the AD brain (Ctrl: 0.16±0.03, AD: 0.3±0.04; p = 0.0282) (Fig 7b, c). Since CHMP2B is a GVD marker and implicated in AD, we also investigated CHMP2B protein levels along with its partner CHMP2A, although we did not see significant difference at the mRNA level. CHMP2B protein showed a significant increase in the AD samples (Fig 7b. c; Ctrl: 0.33±0.09, AD:1.05±0.24; p = 0.0215), whereas no difference was seen for CHMP2A. Co-expression of ESCRT-III components was present in pMLKL+ neurons, with a significant overlap at the subcellular level for CHMP2B, CHMP3 and VPS4B, confirming pMLKL localization in the GVD bodies (Fig 7d). Our data suggest that the activation of necroptosis might alter the expression levels of ESCRT III proteins in AD as a compensatory mechanism.

**Fig. 7.**
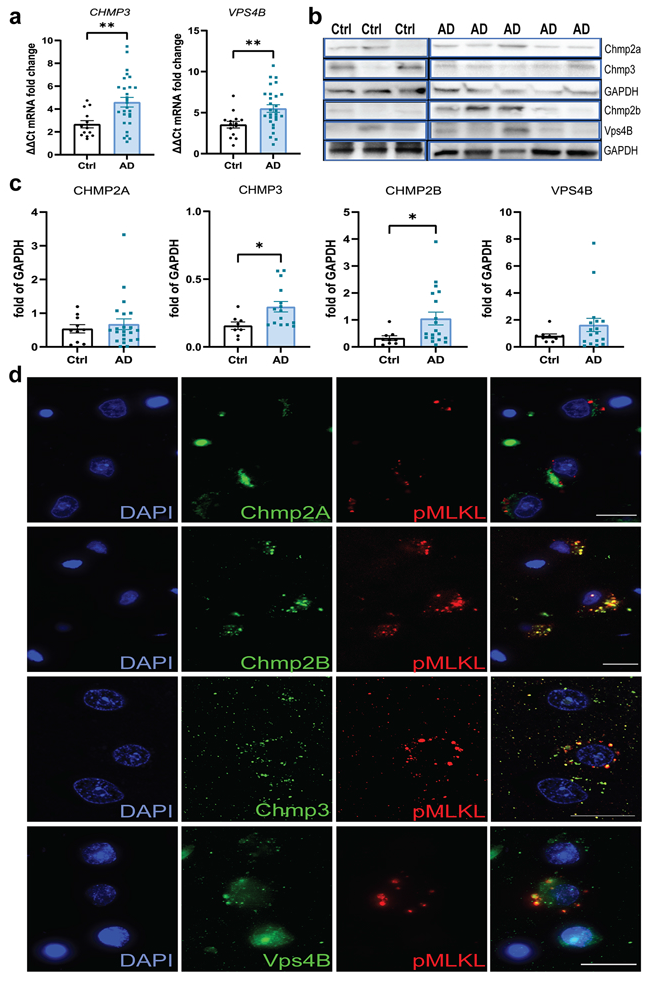
Altered expression of ESCRT III components in the AD hippocampus. **a)** Analysis of mRNA levels of the Vps24, and Vps4b genes in the hippocampus grey matter in AD cases (n=30) and controls (n=14). **b)** Representative western blots of CHMP2A, VPS24, CHMP2B, and VPS4B. Full blots are shown in suppl Fig. 6. **c)** Quantification of CHMP2A, VPS24, CHMP2B, and VPS4B protein levels in AD (n≥10) and control cases (n≥5), normalized to GAPDH. Data are represented as mean ± SEM, **p<0.01. **d)** Representative immunofluorescence image of Chmp2A (green, top row), Chmp2B (green, second row), Vps24 (green, third row), and Vps4B (green, bottom row) with pMLKL (red) and DAPI+ cell nuclei (blue) in the CA1 neurons of AD cases (scale bar = 20μm).

### TNF induces necroptosis activation in iPSC derived human glutamatergic neurons

In order to confirm that TNF can indeed induce necroptosis in human neurons, human IPSC-derived glutaminergic neurons were treated under different conditions for 24 h at day 9-11 post-plating and assessed for cell cytotoxicity by measuring the release of lactate dehydrogenase (LDH) into the culture medium. Before starting the treatments, we ascertained that the neurons exhibited neurite outgrowth and confirmed their homogenous expression of pan-neuronal protein MAP2 and glutamatergic neuron-specific transporter, VGLUT1 (Suppl Fig 4a). TNF alone (100 ng/ml) was not sufficient to induce neuronal cell death, nor were the pan-caspase inhibitor z-Val-Ala-Asp (Ome)-fluoromethylketone (z-VAD; 10 μM) or a SMAC mimetic (SMAC; 2 μM), when added alone (Fig. 8a). TNF and SMAC mimetic together increased LDH release significantly compared to vehicle treatment, which was further increased in a time-dependent manner when all three were combined together (TSZ treatment), reaching 67% cytotoxicity after 24 h treatment (Fig. 8a, Suppl Fig. 4b). To further define the role of necroptosis in cell death induced by TSZ exposure, we pharmacologically inhibited RIPK1, RIPK3, or MLKL with GSK547 (10nM), GSK872 (100nM), or necrosulfonamide (NSA, 1 μM), respectively, and demonstrated that all the inhibitors individually significantly reduced the cytotoxicity by 30-55% (Fig 8b). Increased reactive oxygen species (ROS) production by mitochondria has also been shown to promote RIP autophosphorylation and is required for RIP3 recruitment into the necrosome [66]. Therefore, we determined the levels of intracellular ROS production following TSZ treatment of human neurons, with or without the RIPK1, RIPK3 and MLKL inhibitors. We found that there was a significant increase (>2-fold) in ROS production with TSZ treatment even by 6 h post-treatment, which was effectively reduced by approximately 25% in the presence of the inhibitors (Fig 8c, Suppl Fig 4c).

**Fig. 8.**
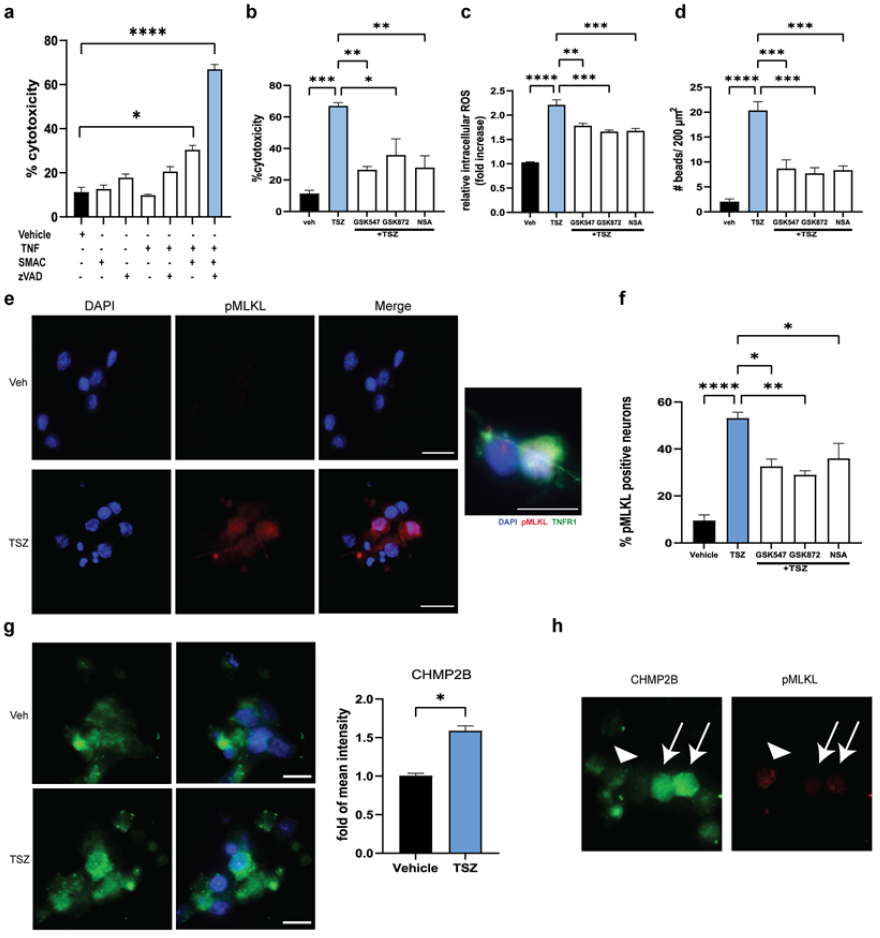
TNF mediated activation of necroptosis in human iPSC-derived glutamatergic neurons when apoptosis is inhibited. **a)** Viability of human iPSC-derived glutamatergic neurons treated with combinations of TNF (100 ng/mL), SMAC mimetic (2μM), and zVAD (10μM) [TSZ], or vehicle control, assessed by LDH assay at 24 h. **b)** Viability of human iPSC-derived glutamatergic neurons treated with vehicle control, TSZ, or TSZ with GSK-547 (10nM), GSK-872 (100nM), or necrosulfonamide (NSA, 1μM), assessed by LDH assay at 24 h. **c)** Quantification of intracellular ROS in human iPSC-derived glutamatergic neurons treated with vehicle control, TSZ, or TSZ with GSK-547, GSK-872, or NSA. **d)** Quantification of neurite beading in human iPSC-derived glutamatergic neurons treated with vehicle control, TSZ, or TSZ with GSK-547, GSK-872, or NSA. **e)** Representative images of human iPSC-derived glutamatergic neurons treated with vehicle or TSZ immunostained for pMLKL (red) and DAPI (blue), along with a representative image of TSZ treated neurons co-stained for pMLKL (red) and TNFR1 (green) (scale bar = 20μm). **f)** Quantification of pMLKL+ neurons treated with vehicle control, TSZ, or TSZ with GSK-547, GSK-872, or NSA. **g)** Representative images of human iPSC-derived glutamatergic neurons treated with vehicle or TSZ immunostained for Chmp2B (green) and DAPI (blue), along with quantification of Chmp2b mean intensity normalized with DAPI mean intensity. **h)** Representative image of Chmp2B (green) and pMLKL (red) expression in human iPSC-derived glutamatergic neurons treated with TSZ. All quantification performed with 3 replicates per group per experiment. One-way ANOVA with Bonferroni’s post-hoc correction for multiple group comparisons, and t-test for comparing between two groups. Data are represented as mean ± SEM, *p<0.05, **p<0.01, ***p<0.001, ****p<0.0001.

Immunostaining with neurofilament-H (NfH) antibody showed axonal/neurite beading at 8 h post-TSZ treatment, which further increased at 16 h post-treatment (Suppl Fig 4d) when compared to vehicle treated neurons. Quantification of the number of beads across different treatment conditions at 16 h of TSZ treatment showed a 10-fold increase compared to vehicle treatment, indicative of neurodegeneration. Treatment with RIPK1, RIPK3 or MLKL inhibitors decreased this number significantly compared to TSZ treatment alone (Fig 8d, Suppl Fig 4d). Next, we investigated the expression of pMLKL as an indication of activation of necroptosis across different treatment conditions at 16 h post-treatment. Upregulation of pMLKL expression was observed in the cytoplasm of the treated neurons in response to TSZ (fig 8e, f). Moreover, pMLKL+ neurons were also positive for TNFR1 expression (Fig 8e). Pre-treatment with RIPK1, RIPK3 and MLKL inhibitors reduced the number of pMLKL positive neurons (Fig 8f).

Since some of the ESCRT III components were altered in the AD brain, we also investigated their expression levels following TSZ treatment in the iPSC-derived glutamatergic neurons. Chmp2B showed a significantly increased expression with TSZ treatment (Fig 8g), with no significant changes observed for Chmp2A, Chmp3, and Vps4B (Suppl Fig. 5). Neurons expressing higher levels of Chmp2B had lower pMLKL expression (arrows) and vice versa (arrowhead) (Fig 8h). Taken together, these results demonstrate that TNF can trigger necroptotic cell death in human glutamatergic neurons through MLKL activation, but only after the alternative TNF-signaling pathways have been inhibited. Small molecule inhibitors of RIPK1, RIPK3, and MLKL were able to significantly reduce cytotoxicity, ROS production and neurite beading in the TSZ treated neurons. In addition, TSZ treatment resulted in increased Chmp2B expression in these neurons, possibly to counteract the increase in pMLKL in those neurons.

## Discussion

Recently, we demonstrated that a local increase in TNFR1 expression on the cortical grey matter neurons in progressive MS can be linked to necroptosis signaling and neurodegeneration, in the presence of reduced apoptotic signaling [42]. Because of the extensive interest in inflammatory mechanisms in neurodegeneration in other neurological disorders [11,15,21,30,52,64], we extended our study to the AD brain. Here, we demonstrate a link between TNF signaling via TNFR1 and necroptosis activation and neuronal loss in the AD hippocampus using human post-mortem brain tissues. This was recapitulated in vitro in human iPSC-derived glutamatergic neurons treated with TNF when apoptosis was inhibited. In addition, our findings show that small molecule inhibitors against RIPK1, RIPK3, and MLKL could significantly protect human cortical neurons against TNF-mediated necroptotic cell death in vivo. We also demonstrate related changes in the ESCRT-III pathway components in the AD hippocampus. These findings provide new insights into the involvement of TNF signaling and necroptosis in neurodegeneration in AD.

Our results showing an increase in the expression of necroptosis pathway genes and proteins in the AD brain, consistent with previous studies on necroptosis activation [5, 28, 41], but build on this to show co-expression of RIPK1 and pRIPK3 with pMLKL in the same neurons, and coimmunoprecipitation of pRIPK3 with MLKL, indicating interaction and necrosome formation in these neurons. Upon phosphorylation, MLKL is suggested to oligomerize in the cytoplasm and migrate to the plasma membrane, where it causes membrane destabilization and ultimately cell death [57]. We also demonstrate the presence of MLKL oligomers, most likely dimers and tetramers, exclusively in the AD samples. Neurons expressing pTau tangles were often also pMLKL positive, suggesting a link between pTau and necroptotic cell death. pMLKL expression levels correlated significantly with pTau expression but not with Aβ expression, consistent with a previous study [28]. However, pMLKL+ neurons were frequently found in close proximity to Aβ plaques. Expression of the activated proteins involved in the final stages of necroptosis, pRIPK3 and pMLKL, was predominantly seen in the large pyramidal neurons of the CA1-2 regions of the hippocampus, which is the most vulnerable region for AD-related neurodegeneration, suggesting that specific neurons in these regions could be differentially vulnerable to the inflammatory microenvironment. While determining the factors that could contribute to a chronic inflammatory microenvironment in the hippocampal region, we observed an increase in activated microglia, assessed by HLA-DR staining and hallmark morphological changes.

Interestingly, the pMLKL+ neurons were often observed in close proximity to these activated microglia with a strong overall correlation between the pMLKL+ neuron density and HLA-DR+ microglial staining. Together with the observation that there was an increase in CD8+ T cells in the perivascular and meningeal regions in the proximity of pMLKL+ neurons in the hippocampus in some AD brains, this demonstrates the presence of neurons with activated necrosome proteins in a potentially inflammatory environment. Recent studies have shown that CD8+ T cells are not only found in the cerebrospinal fluid (CSF) in AD but can also contribute to the progression of cognitive impairment [15, 52], suggesting that these T cells could secrete cytotoxic and proinflammatory cytokines that affect the hippocampus. Since the hippocampus is surrounded by CSF, this creates the possibility for the CD8+ T cells to either infiltrate the hippocampus or for the neurons and microglia to be exposed to the secreted proinflammatory cytokines. Taken together, our data suggest that CA1-2 pyramidal neurons are susceptible to necroptosis under inflammatory conditions, consistent with the hypothesis that inflammation is one of the key drivers of neurodegeneration in AD.

Significantly elevated levels of necroptosis pathway proteins and increased numbers of degenerating neurons have been previously reported in 5xFAD mice, which are characterized by pronounced cell loss, when compared with non-transgenic littermates or with APP/PS1 mice that do not show significant cell loss [5, 41]. These effects were reduced in 5xFAD mice treated with the necroptosis inhibitor, necrostatin (Nec1S), compared to vehicle-treated animals, suggesting that necroptosis contributes to neuronal death in this AD model [5]. Consistent with these observations and previous data from human tissues [28], we observed a strong correlation between the increase in pMLKL+ neuron density and the decrease in overall neuron density in the hippocampus. In addition, we also noted a similar negative relationship between pRIPK3+ neuron density and neuronal loss in the CA1 region. However, whether all neurons expressing these necroptosis activation markers will eventually die, or for how long the neurons express these markers before dying, cannot be ascertained from this data. The percentage of pRIPK3+ and pMLKL+ neurons was relatively high (a mean of 14.5% and 9.5%, respectively) for a single snapshot in time, which suggests that necroptosis is likely to contribute to significant neuronal loss over the disease course. This is significantly higher than that observed previously for MS brains [42]. This difference may reflect the cumulative loss of neurons over a long disease duration in a slowly progressive disease like MS as compared to a more rapid neuron loss in a late-onset condition with a shorter disease duration like AD. Interestingly, we observed a significantly higher number of pRIPK3+ and pMLKL+ neurons in the female AD brains compared to male AD brains. Taken along with our data showing a strong negative correlation between neuron density and pMLKL/pRIPK3 positive neurons, this could translate to increased neuron loss in the female AD brain than the male AD brain over the disease course. This supports previous findings that in patients with mild cognitive impairment and AD, brain volumes decline faster in women than in men [50], and that a negative correlation between Braak stages and neuronal density in hippocampus was observed in women, specifically in the CA1 region [32].

Apoptosis was not significantly activated in the AD brain compared to the control brain, supporting previous studies that show that caspase inhibition or insufficient caspase activation can lead to necroptosis [1, 13, 19]. Our observations showed that there was absence of apoptotic morphological features, and no caspase 8 activation in the AD and control brains. The absence of caspase 8 activation not only leaves the RIPK1 kinase domain intact to autophosphorylate and then phosphorylate downstream RIPK3 and MLKL [26, 57], but has also been shown to disturb the inflammatory homeostasis in mice [29]. Moreover, two Caspase 8 variants have been reported to be associated with AD in a large cohort study, both of which showed reduce functionality due to either significantly lower activity or sequestration [45]. Activated caspase 3 levels were lower in the AD hippocampus compared to the control hippocampus. In addition, at the mRNA level, caspase 3, and cIAP1 levels were decreased in the AD hippocampal tissue, suggesting reduced apoptosis in the AD tissues. The reduced *CFLAR* mRNA levels suggest decreased NF-κB activation, which would also increase necroptotic signaling in the absence of apoptosis [7, 49, 63]. Taken together with our data on the necroptosis pathway activation, these results point to necroptosis as a major form of cell death in the AD brain, although other forms of cell death, such as ferroptosis and pyroptosis cannot yet be excluded.

Although necroptosis activation has been recently demonstrated in AD [5, 28], the mechanism underlying the activation remains unclear. We propose that pro-inflammatory cytokines, such as TNF, can diffuse into the hippocampal grey matter from the CSF or be induced in glial cells by dying neurons/Aβ plaques/reactive microglia, to induce necroptosis and neurodegeneration via TNFR1 signaling, similar to our findings in MS cortical grey matter [6, 31, 42]. The increase in TNF levels in the hippocampal parenchyma and the co-expression of pRIPK3 and pMLKL in neurons with increased TNFR1 expression strongly suggests the activation of TNFR1-mediated signaling pathway in the hippocampal neurons in AD. Although TNFR1 signaling can trigger apoptosis, a decrease or no change in caspase-8 and caspase-3 cleavage, along with the increase in pRIPK3 and pMLKL levels, indicates a switch towards necroptosis rather than apoptosis in these neurons. In the presence of a caspase inhibitor and a SMAC mimetic, which extrinsically recapitulates our observations in the AD brain, TNF stimulation of iPSC derived human glutaminergic neurons led to the activation of necroptosis by increasing the pMLKL expression in the cytoplasm, which was reversed by RIPK1, RIPK3, or MLKL inhibitors. Interestingly, these inhibitors not only reduced the pMLKL expression in the treated neurons, but also largely restricted the pMLKL expression to the nucleus. This suggests that the presence of cytosolic pMLKL is important for necroptotic cell death, consistent with the finding that blocking nuclear export and increasing nuclear retention of pMLKL, reduces necroptotic cell death [60].

Using a rat model in which lentiviral vectors carrying the transgenes for human TNF and interferon (IFN)-γ were injected into the subarachnoid space to induce persistent cytokine production by meningeal cells, we have previously shown that persistently increased TNF and IFNγ in the CSF is sufficient to induce neuronal loss along with upregulation of TNFR1 and pMLKL in the areas with extensive inflammation [42]. This further supports our hypothesis that TNF-mediated inflammation, be it in vitro, in vivo, or during pathological conditions within the human brain, is sufficient to induce cell death through necroptosis activation.

In addition to the direct stimulation of necroptosis by TNF, this pro-inflammatory cytokine may play a role in neurodegeneration by stimulating ROS generation in neurons and glia [11, 40, 46] and promoting glutamate release by microglia [14, 51]. In addition to the reduction in cell death and pMLKL expression, pre-treatment with RIPK1, RIPK3 and MLKL small molecule inhibitors also resulted in reduced beading of neurites, indicative of reduced neurodegeneration, and reduced intracellular ROS caused by TNF signaling. The latter is of significance as studies have shown that ROS functions in a positive feedback loop to ensure effective necroptosis activation, especially in the presence of TNF and a SMAC mimetic [47]. TNF can induce mitochondrial ROS, although ROS involvement in necroptosis is suggested to be cell context dependent [17, 20]. ROS can activate RIPK1 autophosphorylation and is essential for RIPK3 recruitment into the necrosome, thus enhancing necrosome formation. Conversely, ROS induction requires necrosomal RIPK3 [66]. Our data supports the role of ROS as a critical regulator of necroptosis in the context of TSZ treatment and the significant reduction of intracellular ROS by pretreatment with necroptosis inhibitors correlates with increased cell survival.

ESCRT-III proteins, which are involved in endosomal trafficking, virus budding, and multivesicular body formation [43], have been shown to act downstream of MLKL, antagonising necroptosis stimulated by TNF in vitro by extending the duration of plasma membrane integrity and sustaining cell survival [16]. Among the ESCRT-III components, mutations in the Chmp2B gene are reported to be associated with frontotemporal dementia (FTD) and amyotrophic lateral sclerosis (ALS) [56], which provides a clear link between dysfunction in the ESCRT-III machinery and neurodegeneration. Moreover, Chmp2B has been considered to be a GVD marker, as it shows strong immunoreactivity in the GVD bodies in AD brain [61]. However, to date there have been no reported associations between AD and known mutations of any ESCRT-III pathway components. Our data showed an increase in Chmp2B expression in the AD brain, specifically in the GVD bodies, and also in human iPSC-derived glutamatergic neurons treated with TSZ. Chmp2B expression also colocalized with pMLKL expression in the AD brain, thus confirming the presence of pMLKL in the GVD bodies. Although most studies have focused on the effects of overexpression of mutated Chmp2B (truncated) relevant to FTS and ALS, one study showed that overexpression of full-length Chmp2B in HeLa cells caused deformation of the plasma membrane [3]. Among other components, there was an increase in the expression of Vps24/Chmp3 in the AD brain, but not in cultured neurons. Chmp3 is required for the protein sorting and formation of multivesicular bodies (MVB) [2]. A constant recycling of Chmp3 by Vps4 is required to promote the net growth of ESCRT-III assemblies [33], and an overall increase in the levels of Chmp3 in the AD brain might indicate decreased turnover of Chmp3. A recent study showed that MAPT/Tau accumulation disrupts ESCRT-III complex formation in mice, thus impairing lysosomal degradation [10], which might explain the link between pTau expression and necroptosis in neurons, as shown in our study and others [28]. It has also been reported that MLKL and pMLKL levels can affect endosomal transport independently of RIPK3 and are required for the generation of intraluminal and extracellular vesicles [62]. Thus, the lack of effective extrusion of pMLKL from the plasma membrane into endosomes or exosomes may be responsible for vulnerability of certain cells to necroptosis. However, given the higher-than-expected number of neurons in the AD brain showing necroptosis markers along with ESCRT-III components, and the heterogeneity in the number of pMLKL positive puncta across these neurons, it could mean that they are trying to overcome necroptosis and some will survive the insult. A subset of them may then fail to overcome the necroptotic signaling in the presence of additional factors such as pTau or proximity to higher levels of inflammatory mediators. Therefore, whether the increase in Chmp2B and Vps24 expression in the AD brain is the result of a compensatory mechanism that has been triggered in response to necroptosis to help neurons survive or is a dysregulation of their function that then results in failure to protect the neurons from necroptotic cell death is unclear. Detailed studies will be required to establish functional link between necroptosis and the ESCRT III pathway in neurons in vivo under pathological conditions.

In conclusion, we provide strong evidence for inflammation-driven activation of TNF-mediated necroptosis in CA1-2 hippocampal neurons in the post-mortem AD brain and propose that targeting the necroptosis pathway could hold great promise for inhibiting neurodegeneration. Our data suggests that certain neurons are more susceptible to necroptotic cell death, and that this selective neuronal vulnerability may be due to an inflammatory microenvironment contributed by activated microglia or CD8+ T cells that secrete proinflammatory cytokines, or due to dysregulation/imbalance in the ESCRT-III pathway or increased co-expression of pTau within those neurons. While our results indicate that TNF-TNFR1 interaction may be a key mechanism that drives neurodegeneration in AD, it is important to note that there are other triggers and pathways leading to necroptotic cell death such as FasL, TRAIL, dsRNA, and viral Z-RNA [27, 37, 65], although their role in neuronal loss in AD is yet to be investigated. Emerging evidence of necroptosis in several neurodegenerative diseases and our own studies in MS and AD suggests convergence of molecular mechanisms underlying neuronal loss in different pathologies. Identifying the precise mechanisms underlying the loss of neurons in AD is important for developing effective strategies for the treatment of AD. Activation of RIPK1 has been demonstrated to mediate the majority of the pro-inflammatory and cell death inducing signaling stimulated by TNF/TNFR1 interaction, including those in the CNS [63]. RIPK1 is a key mediator of cell death and small molecule inhibitors of RIPK1 have advanced into clinical trials for several CNS and non-CNS disorders [18, 34]. Using small molecule inhibitors against RIPK1 and or other necroptosis pathway components in a suitable chronic animal model for AD showing neuronal loss would, therefore, provide definitive proof of a role for TNF/TNFR1 stimulated necroptosis in AD.

## Author contributions

AJ contributed to concept and design, acquisition, analysis, and interpretation of data and drafting of manuscript. TTH and RJ contributed to acquisition of data. CP contributed to concept and design. RR contributed to concept and design, obtaining research grants, interpretation of data and revision of the manuscript.

## Declaration of interests

AJ, TTH, RJ, CP and RR have no competing interests concerning this study.

## Acknowledgments

We thank the UK Multiple Sclerosis and Parkinson’s Tissue Bank at Imperial College London (funding from the MS Society of Great Britain, grant 007/14 to RR) and South West Dementia Brain Bank, University of Bristol, for the supply of post-mortem AD and control brain tissue samples. We thank Anselm Vincent for his assistance with technical support. This work was supported by a LKCMedicine Strategic Academic Initiative to RR.

## Notes

### Competing Interest Statement

The authors have declared no competing interest.

