## Supplemental figures and tables for "TNF-mediated neuroinflammation is linked to neuronal necroptosis in Alzheimer’s disease hippocampus"

Supplementary Table 1: Cohort characteristics

| Case number | Age (years) | Sex | Braak stage[4] | CERAD score[35] | Clinical diagnosis | Post-mortem delay (h) |
|---|---|---|---|---|---|---|
| 1 | 85 | M | 2 | 1 | Non-AD control | 30.5 |
| 2 | 74 | F | 1 | 0 | Non-AD control | 39.5 |
| 3 | 83 | M | 3 | 0 | Non-AD control, mild CAA | 37.5 |
| 4 | 78 | M | 2 | 0 | Non-AD control | 56.25 |
| 5 | 95 | F | 2 | 0 | Non-AD control | 59.25 |
| 6 | 83 | M | 2 | 0 | Non-AD control | 47.75 |
| 7 | 89 | F | 2 | 0 | Non-AD control | 26.5 |
| 8 | 93 | M | 2 | 0 | Non-AD control | 34.5 |
| 9 | 78 | F | 2 | 0 | Non-AD control | 66.25 |
| 10 | 92 | M | 2 | 0 | Non-AD control, CAA | 47.25 |
| 11 | 63 | F | 1 | 0 | Non-AD control | 21 |
| 12 | 66 | F | 6 | 4 | AD, CAA, Dementia | 37.25 |
| 13 | 93 | F | 5 | 4 | AD | 31.75 |
| 14 | 91 | F | 5 | 3 | AD, Dementia | 28.25 |
| 15 | 87 | F | 5 | 3 | AD | 40.5 |
| 16 | 73 | F | 6 | 3 | AD | 25.75 |
| 17 | 86 | M | 4 | 3 | AD, HS | 34.5 |
| 18 | 88 | F | 5 | 3 | AD, SVD, Dementia | 80.5 |
| 19 | 75 | F | 5 | 3 | AD, BA, VaD | 37.5 |
| 20 | 80 | M | 6 | 3 | AD, Dementia | 21.25 |
| 21 | 57 | F | 6 | 3 | AD, CAA, Early-onset Dementia | 30.5 |
| 22 | 89 | F | 6 | 3 | AD | 31.25 |
| 23 | 80 | M | 5 | 2 | AD | 43 |
| 24 | 92 | F | 6 | 3 | AD | 39.5 |
| 25 | 89 | F | 5 | 3 | AD, SVD, Dementia | 35.75 |
| 26 | 86 | F | 5 | 3 | AD | 37.5 |
| 27 | 83 | M | 3 | 1 | AD, MVD | 45.5 |
| 28 | 85 | F | 6 | 3 | AD. CAA | 17.75 |

| | | | | | | |
|---|---|---|---|---|---|---|
| 29 | 86 | M | 5 | 3 | AD, VaD | 47.5 |
| 30 | 83 | M | 3 | 3 | AD, Dementia | 41.75 |
| 31 | 99 | F | 5 | 3 | AD, SVD, Dementia | 43 |
| 32 | 90 | M | 5 | 3 | AD | 61 |
| 33 | 78 | F | 6 | 3 | AD, Dementia | 27.75 |
| 34 | 87 | M | 6 | 3 | AD, CAA, SVD, Dementia | 44.25 |
| 35 | 70 | M | 6 | 3 | AD, Dementia | 57 |
| 36 | 84 | F | 4 | 1 | AD | 22 |
| 37 | 85 | M | 4 | 2 | AD | 23 |
| 38 | 68 | M | 4 | 2 | AD | 24 |
| 39 | 84 | F | 4 | 3 | AD, Dementia | 13.5 |
| 40 | 80 | F | 4 | 2 | AD | 28 |
| 41 | 87 | M | 5 | 3 | AD | 41.75 |

*AD* Alzheimer's disease, *F* female, *M* male, *CAA* cerebrovascular angiopathy, *HS* hippocampal sclerosis, *SVD* subcortical vascular dementia, *BA* basal atherosclerosis, *MVD* moderate vascular disease, *VaD* vascular dementia. CERAD scores for neuritic plaque densities: 0 = none (no plaques), 1 = sparse, 2 = moderate, and 3 = High density.

Supplementary Table 2: Antibody list

| REAGENTS | SOURCE | IDENTIFIER | APPLICATION |
|---|---|---|---|
| Mouse monoclonal anti-human PHF-Tau antibody | Thermo Scientific | Cat# MN1020 **RRID:AB_223647** | IF and IHC (1:500) |
| Mouse monoclonal anti-amyloid βA4 (1E8) | Millipore | Cat# MABN639 | IF and IHC (1:500) |
| Mouse monoclonal anti-GAPDH antibody | Bio-Rad | Cat#VMA00046 | WB (1:1000) |
| Mouse monoclonal anti-MLKL (32B) antibody | Santa Cruz Biotechnology | Cat#sc-293201 | WB (1:1000) |
| Rabbit polyclonal anti-phospho-human MLKL (Ser 358) antibody | Abcam | Cat# ab187091 **RRID:AB_2619685** | WB (1:1000); IHC and IF (1:200) |
| Mouse monoclonal anti- human RIPK3 antibody | R&D Systems | Cat# MAB7604 **RRID:AB_2619684** | WB (1:1000) |
| Rabbit monoclonal anti-phospho-human RIPK3 (Ser 227) antibody | Cell Signaling Technology | Cat# 93654 **RRID:AB_847259** | IHC and IF (1:250) WB (1:1000) |
| Mouse monoclonal anti-RIPK1 (7H10) antibody | Abcam | Cat# ab72139 **RRID:AB_2178115** | WB (1:1000); IF (1:200) |
| Rabbit monoclonal anti-TNF receptor II antibody | Abcam | Cat#ab109322 | WB (1:1000) |
| Mouse monoclonal anti-NeuN antibody | Millipore | Cat# MAB377 | IF (1:1000) |
| Rabbit monoclonal anti-FADD antibody | Abcam | Cat# ab108601 **RRID:AB_10864812** | WB (1:1000) |
| Rabbit polyclonal anti-active caspase 3 antibody | R&D systems | Cat# AF835 **RRID:AB_2243952** | WB (1:1000) IF (1:200) |
| Rabbit monoclonal anti-CD8 (SP16) antibody | Invitrogen | Cat# MA5-14548 **RRID:AB_10984334** | IHC (1:250) |
| Mouse monoclonal anti-caspase-8 (IC12) antibody | Cell Signaling Technology | Cat# 9746 | WB (1:1000) |
| Mouse monoclonal anti-HuC/D antibody (16A11) | Invitrogen | Cat# A-21271 **RRID:AB_221448** | IHC (1:500) |
| Mouse monoclonal anti-human TNFRI antibody | R and D Systems | Cat# MAB225 **RRID:AB_2204150** | WB (1:1000) |
| Mouse monoclonal anti-Neurofilament 200 kDa (RT97) | Millipore | Cat# MAB5262 | IF (1:1000) |
| Rabbit monoclonal anti-TNF alpha antibody | Abcam | Cat# ab215188 | WB (1:1000) |
| Rabbit polyclonal anti-TNFR1 antibody | Abcam | Cat# ab223352 | IF (1:100) |
| Rabbit polyclonal anti-VPS4B/MIG1 antibody | Abcam | Cat# ab224736 | WB (1:1000) IF (1:200) |
| Mouse monoclonal anti-MAP2 antibody | Abcam | Cat# ab254143 | IF (1:50) |
| Rabbit monoclonal anti-VGlut1 antibody | Abcam | Cat# ab227805 **RRID:AB_2868428** | IF (1:500) |
| Rabbit monoclonal anti-VPS24 antibody | Abcam | Cat# ab175930 | WB (1:1000) |

| | | | |
|---|---|---|---|
| Rabbit monoclonal anti-CHMP2B antibody | Abcam | Cat# ab157208 **RRID:AB_2885096** | WB (1:1000) IF (1:250) |
| Mouse monoclonal anti-human HLA-DR/DP/DQ (CR3/43) antibody | Thermo Scientific | Cat# MA1-25914 **RRID:AB_794857** | IHC (1:500) |
| Rabbit polyclonal anti-GFAP antibody | Abcam | Cat# ab7260 **RRID:AB_305808** | IHC (1:500) |
| ImmPACT® DAB Substrate, Peroxidase (HRP) | Vector Laboratories | Cat# SK-4105 | |
| Vector® Blue Substrate Kit, Alkaline Phosphatase (AP) | Vector Laboratories | Cat# SK-5300 | |
| VECTASTAIN® ABC-AP Kit, Alkaline Phosphatase | Vector Laboratories | AK-500 | |
| ImmPRESS® HRP Horse Anti-Rabbit IgG Polymer Detection Kit, Peroxidase | Vector Laboratories | Cat# MP-7401, | |
| ImmPRESS® HRP Horse Anti-Mouse IgG Polymer Detection Kit, Peroxidase | Vector Laboratories | Cat# MP-7422, | |
| Goat anti-Rabbit IgG (H+L) Secondary Antibody, Alexa Fluor 488, preabsorbed | Abcam | Cat# ab150081 **RRID:AB_2734747** | IHC (1:5000) |
| Goat Anti-Mouse IgG (H+L) Antibody, Alexa Fluor 488, preabsorbed | Abcam | Cat# ab150117 **RRID:AB_2688012** | IHC (1:5000) |
| Goat anti-Mouse IgG (H+L) Secondary Antibody, Alexa Fluor 647, preabsorbed | Abcam | Cat# ab150119 **RRID:AB_2811129** | IF (1:2000) |
| Goat anti-Rabbit IgG (H+L) Secondary Antibody, Alexa Fluor 594, preabsorbed | Abcam | Cat# ab150084 **RRID:AB_2734147** | IF (1:2000) |
| Goat anti-Rabbit IgG (H+L) Secondary Antibody, Alexa Fluor 555, preabsorbed | Abcam | Cat# ab150086 **RRID:AB_2890032** | IHC (1:5000) |
| Goat anti-Mouse IgG (H+L) Secondary Antibody, Alexa Fluor 555, preabsorbed | Abcam | Cat# ab150118 **RRID:AB_2714033** | IHC (1:5000) |
| Goat anti-Mouse IgG (H/L) HRP | Bio-Rad | Cat# 0300-0108P | WB (1:20000) |
| Goat anti-Rabbit IgG HRP | Bio-Rad | Cat# 403005 | WB (1:20000) |

Supplemental Fig 1

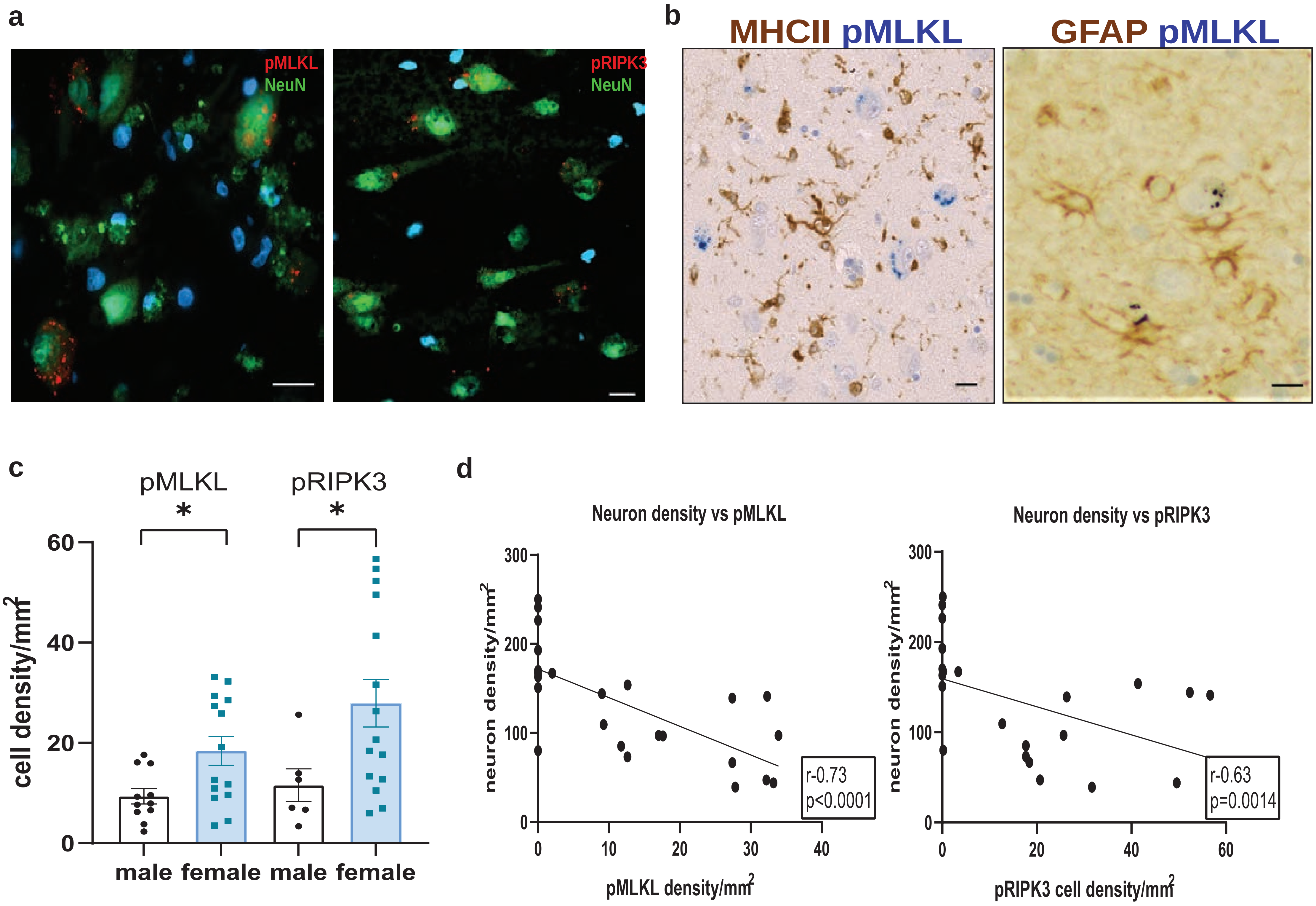


**Supplemental Fig. 1 a)** Representative images showing NeuN+ (green) neuron-specific expression of pMLKL (red, left) and pRIPK3 (red, right) (scale bar = 20µm). **b)** Representative images showing no overlap between microglia (MHC II+, brown) or astrocytes (GFAP+, brown) with pMLKL+ neuronal staining in the CA1 hippocampus of AD brain. **c)** Quantification showing sexual dimorphism in the pMLKL+ and pRIPK3+ cell density in the AD cases. Data are represented as mean ± SEM, *p<0.05. **d)** Correlation analyses between neuron density and pMLKL+ or pRIPK3+ neuron density in AD and control cases. Correlation analysis by Spearman comparison.

Supplemental Fig 2

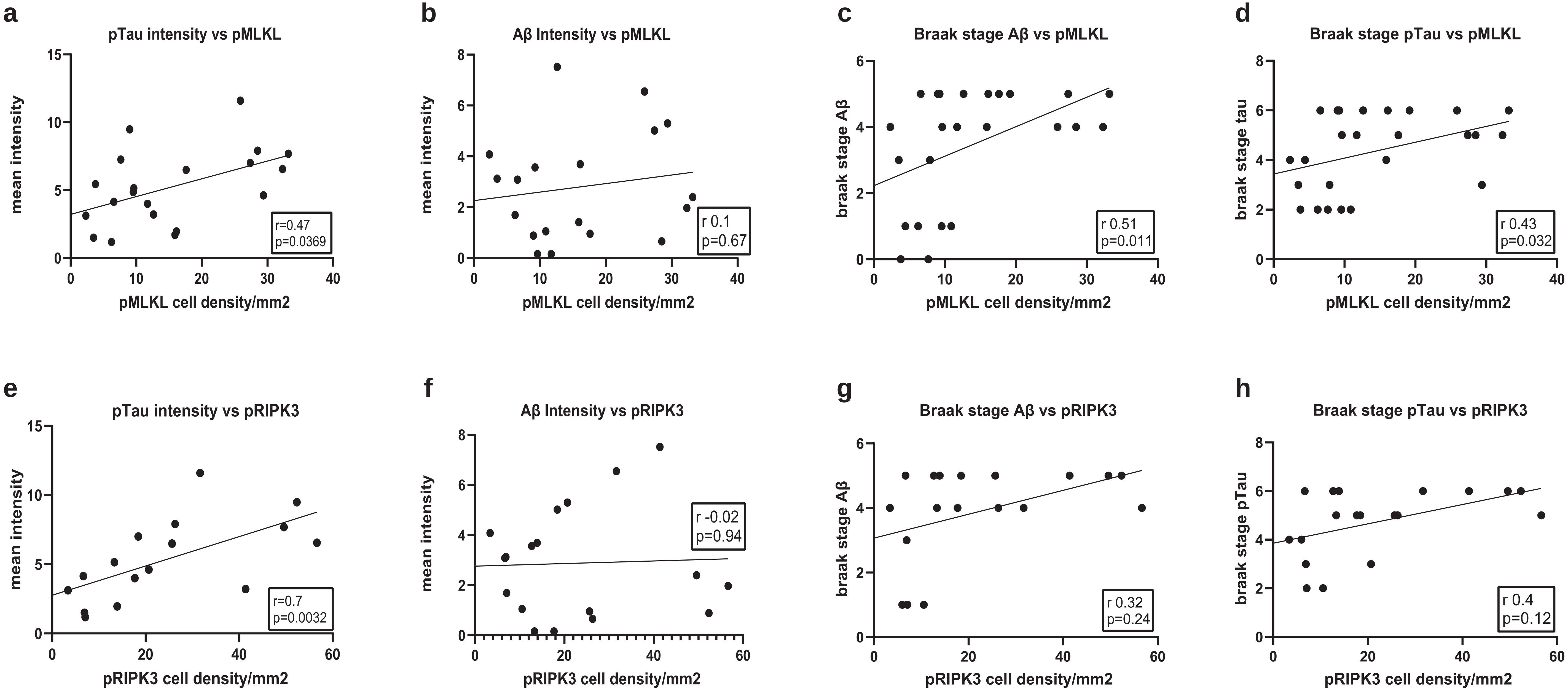


**Supplemental Fig. 2** Association between pRIPK3 cell density and pMLKL cell density with AD pathology. Correlation analyses showing the relationship between pRIPK3 and pMLKL cell densities with pTau and Aβ Braak stages and mean intensities in the AD hippocampus (n≥18). Correlation analysis by Spearman comparison.

Supplemental Figure 3

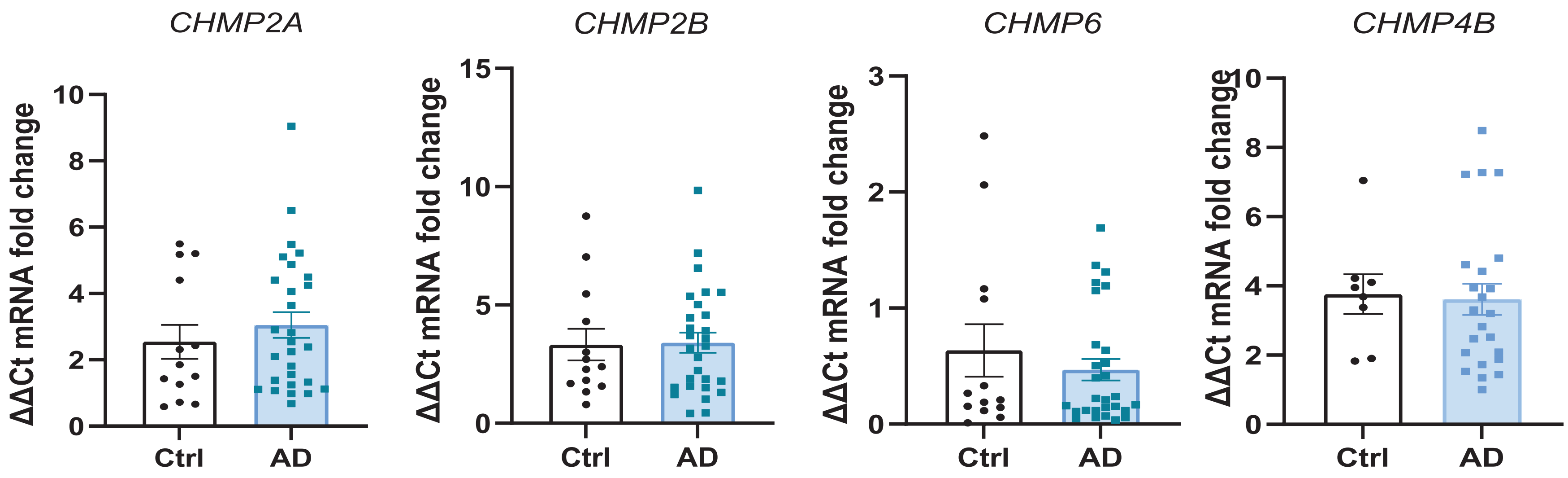


**Supplemental Fig. 3** Analysis of mRNA levels of the chmp2a, chmp2b, chmp6, and chmp4b genes in the hippocampus grey matter in AD cases (n=30) and controls (n>8).

Supplemental Fig 4

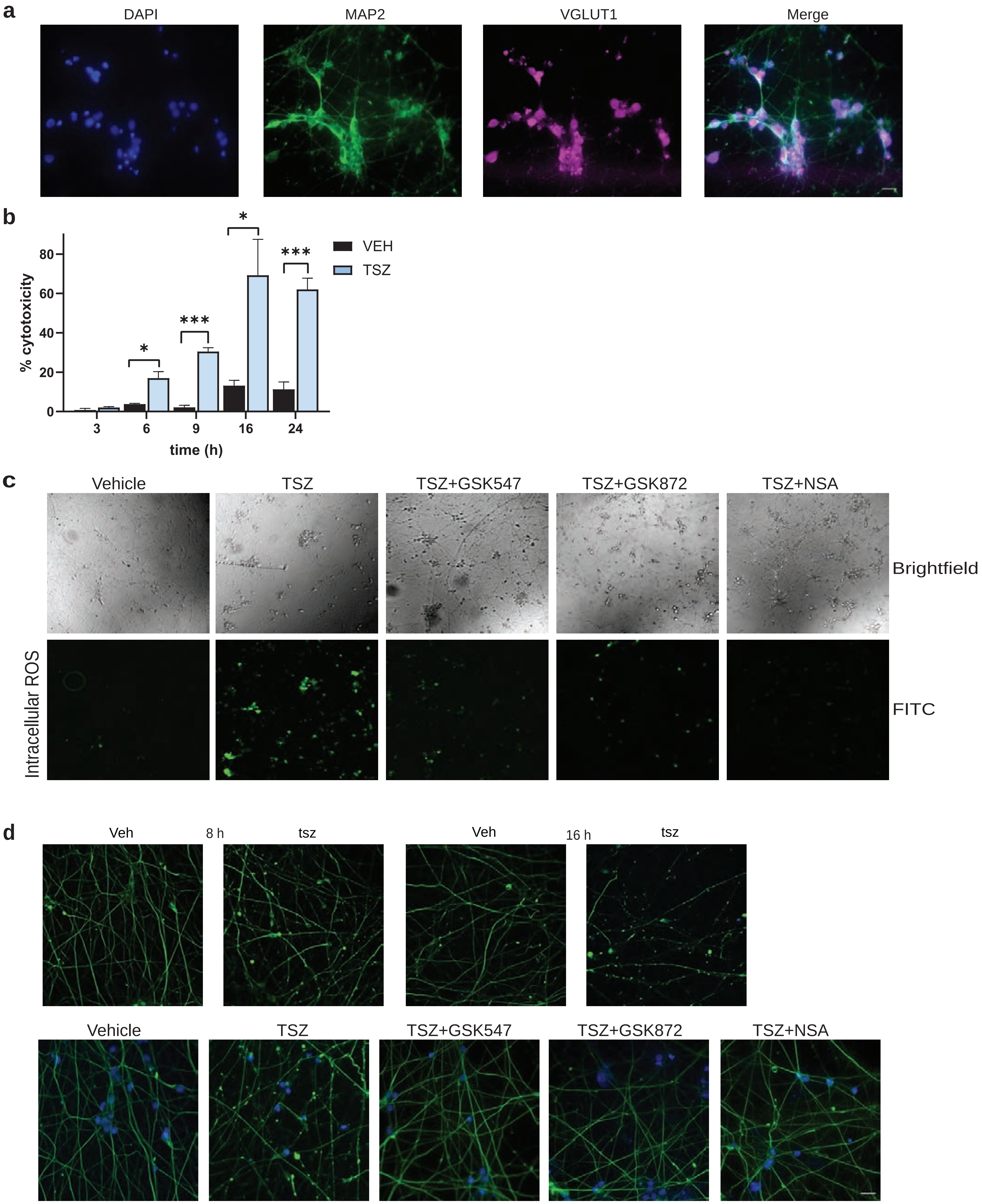


**Supplemental Fig. 4** Characterization of human iPSC-derived glutamatergic neurons under different treatment conditions. **a)** Representative images of MAP2 an VGlut1 expression in the iPSC culture wells. **b)** Cytotoxicity of neurons treated with TNF (100ng/mL), SMAC mimetic (2μM) and zVAD (10μM) at different time points, assessed by measuring LDH release. **c)** Representative brightfield and fluorescence images of iPSC-derived glutamatergic neurons showing intracellular ROS expression under different treatment conditions at 6 h post-treatment. **d)** Top panels: Representative images of NfH in neurons treated with vehicle and TSZ after 8 h and 16 h (scale bar = 20μm), showing beading response with TSZ treatment at both time points. Bottom panels: Representative images of NfH in neurons treated under different conditions at 16 h post-treatment (scale bar = 20μm).

Supplemental Fig 5

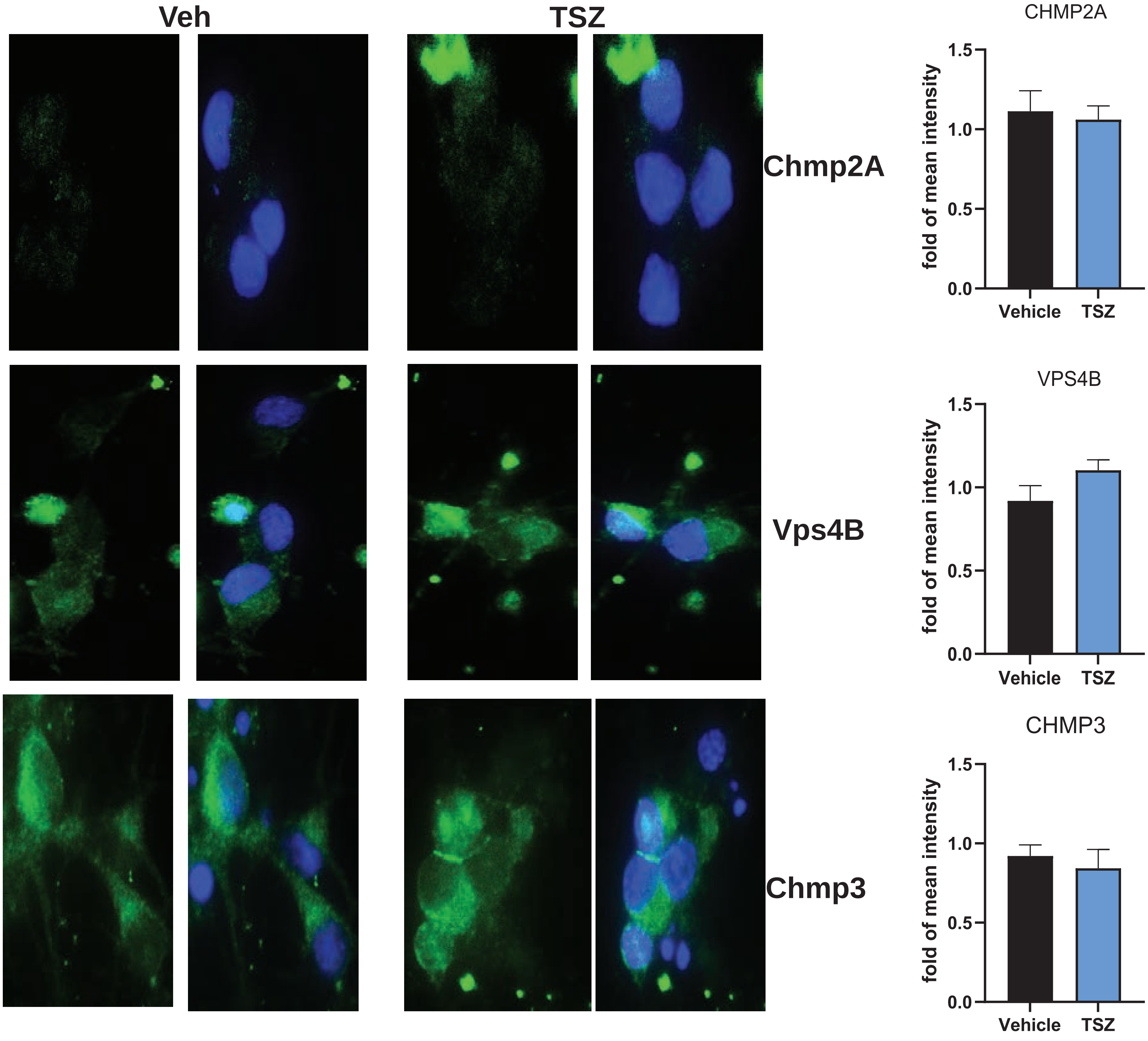


**Supplemental Fig. 5** Expression of ESCRT III components and TNFR1 in response to TSZ. Representative images and quantification of Chmp2a, Vps4b, and Chmp3 with vehicle and TSZ treatment after 16 h. Paired t-tests were performed to compare between the two groups. Data are represented as mean ± SEM. Data are represented as mean ± SEM.

Supplemental Fig 6

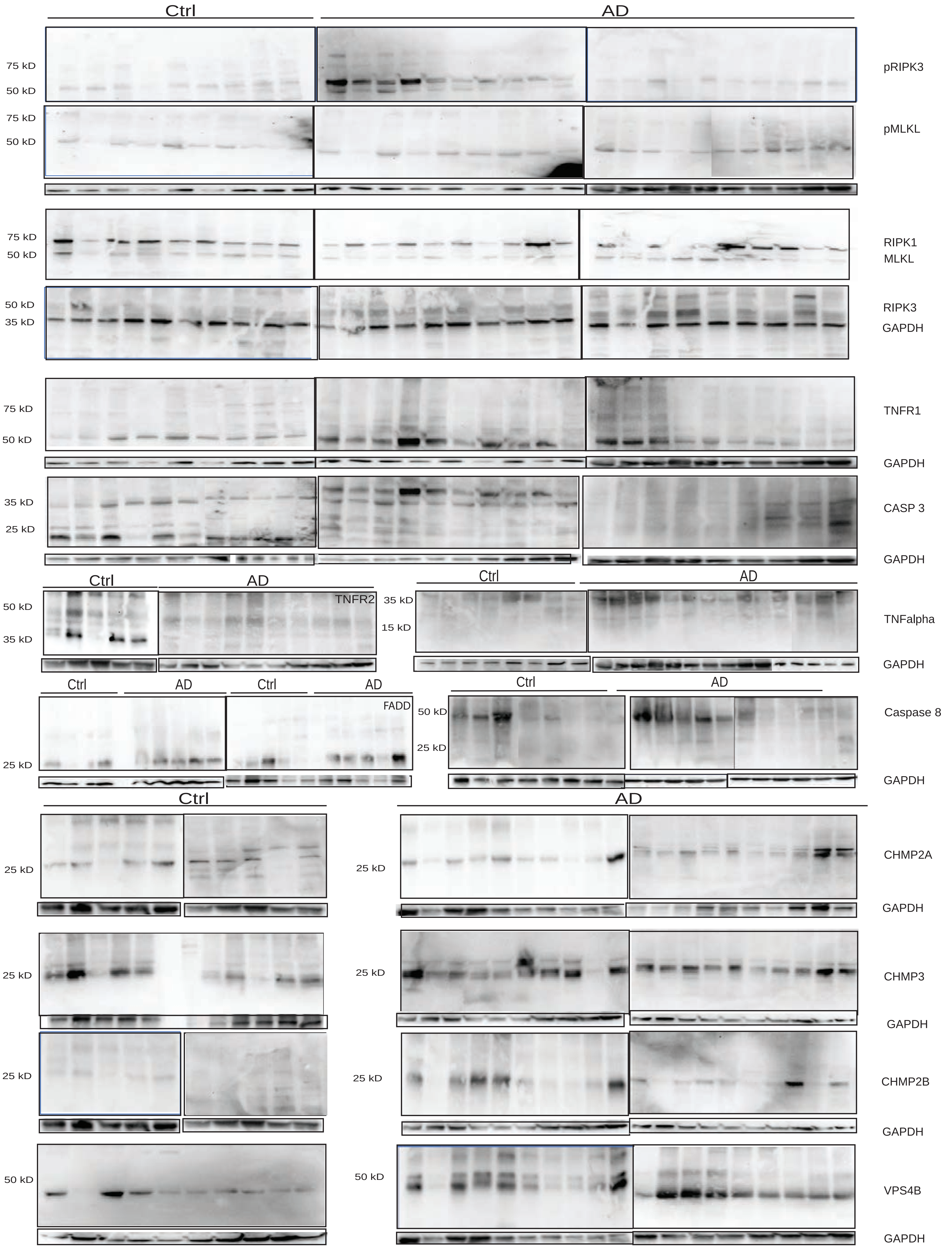


**Supplemental Fig. 6** Full blots of pRIPK3, pMLKL, RIPK1, MLKL, RIPK3, TNFalpha, TNFR1, TNFR2, FADD, Caspase 3, Caspase 8, Chmp2A, Chmp2B, Chmp3, and Vps4B from samples from human post-mortem brains from AD and control cases. Proteins were extracted with RIPA, except for RIPK1, MLKL, and RIPK3, which were extracted with RIPA, followed by UREA. All protein levels were normalized to GAPDH.
